# De novo design of protein binders that target DELE1 to inhibit the mitochondrial stress response

**DOI:** 10.64898/2025.12.22.695711

**Authors:** Rui Yang, Kaiyuan Zheng, McGuire Metts, Yiluo Wang, Danyan Yin, Kevin P. Li, Agnieszka A. Prazmowska, David F. Kashatus, Brian Kuhlman, Jie Yang

**Author notes:** Correspondence: Jie Yang. These authors contributed equally (Rui Yang, Kaiyuan Zheng).

## Abstract

Mitochondrial stress activates the integrated stress response (ISR) through the mitochondrial protein DELE1, which relays stress signals to the cytosolic kinase HRI to induce ATF4. Dysregulation of DELE1-mediated signaling has been implicated in pathological conditions, yet molecular strategies to modulate DELE1 remain unavailable. Here, we report de novo designed proteins that bind DELE1, block its oligomerization, and inhibit DELE1-mediated ISR activation. Several designs form stable complexes with DELE1 and disrupt its oligomerization in vitro while preserving DELE1’s ability to bind HRI. In cells, these designs suppress ATF4 induction during mitochondrial stress and impair the recovery of elongated mitochondrial morphology following transient insult. Crystal structure of a representative binder, together with structural modeling and targeted mutagenesis, confirm that the designed proteins engage a critical interface required for DELE1 oligomerization. These findings establish DELE1 as a druggable target and demonstrate that de novo designed proteins offer precise tools to modulate this pathway, providing a foundation for future therapeutic exploration.

**SIGNIFICANCE:** Mitochondrial stress activates the integrated stress response through the signaling protein DELE1, but no molecular tools have been available to directly target DELE1 and selectively modulate this pathway. We developed de novo designed protein binders that recognize a critical oligomerization interface in DELE1, disrupt its assembly, and suppress mitochondrial stress-induced ISR activation in cells. These binders also impair recovery of mitochondrial network morphology following transient stress, linking DELE1 assembly to adaptive remodeling. Our study establishes DELE1 as a tractable and druggable target in mitochondrial stress signaling and demonstrates that de novo protein design can generate precise modulators of intracellular stress-response pathways.

## INTRODUCTION

Mitochondria are essential organelles best known for their role in oxidative phosphorylation, but they also support numerous processes that influence nearly every aspect of eukaryotic cell physiology. Beyond ATP production, mitochondria function as dynamic signaling hubs that coordinate cellular stress responses, maintain metabolic balance and proteostasis, and regulate nuclear gene expression through retrograde communication^1^. Disruption of mitochondrial function contributes to diverse human diseases, including neurodegeneration^2–5^, metabolic disorders^6–8^, and cancer^9–11^. In mammalian cells, perturbations that impair mitochondrial activity or disturb mitochondrial proteostasis generate mitochondrial stress, which activates the integrated stress response (ISR)^12,13^. The ISR is initiated by four stress-sensing kinases, including HRI, PERK, PKR, and GCN2, that detect distinct upstream insults yet converge to phosphorylate the α subunit of eukaryotic initiation factor 2 (eIF2α)^14^. Phosphorylated eIF2α reduces formation of the translational ternary complex, resulting in global attenuation of translation initiation while selectively enhancing translation of stress-responsive mRNAs. Among these, activating transcription factor 4 (ATF4) functions as the central effector of the ISR, inducing gene-expression programs that restore amino acid balance, redox homeostasis, and proteostasis^15–18^. During prolonged or unresolvable stress, sustained ATF4 activity instead promotes apoptotic cell death.

Recent studies identified the mitochondrially targeted protein DELE1 as an essential link between mitochondrial stress and ISR activation^19,20^. Under resting conditions, full-length DELE1 is imported into mitochondria and rapidly degraded, preventing inappropriate ISR induction^21^. Mitochondrial stress activates the inner mitochondrial membrane protease OMA1, which cleaves DELE1 within the intermembrane space to generate a stable C-terminal fragment, hereafter referred to as DELE1^CTD^, that accumulates in the cytosol. Cytosolic DELE1^CTD^ engages and activates the HRI kinase, leading to eIF2α phosphorylation and ATF4 induction. We recently determined the high-resolution cryo-EM structure of DELE1^CTD^, revealing its oligomeric architecture, and showed that DELE1 oligomerization promotes HRI activation and amplifies downstream ISR signaling^22^.

Despite the central role of DELE1 in mitochondrial surveillance and its increasing implications in human disease^23–31^, no molecular tools or inhibitors exist that directly modulate DELE1 itself. Such DELE1 modulators would enable mechanistic interrogation of the OMA1-DELE1–HRI-ATF4 pathway and provide a foundation for evaluating the therapeutic potential of targeting DELE1 in pathological contexts associated with mitochondrial stress. In this study, we applied a de novo protein design pipeline to generate designed proteins that selectively target the first α-helix of the tetratricopeptide repeat domain within DELE1^CTD^. We screened and validated designs that bind DELE1, form stable complexes in vitro, disrupt its oligomerization, and attenuate ATF4 induction under distinct mitochondrial stress conditions. The crystal structure of a representative design, together with structural modeling and targeted mutagenesis, confirms that the designed DEEL1 binders engage a critical interface required for DELE1 self-assembly, thereby blocking formation of the active oligomeric scaffold. Together, these findings establish DELE1 as a druggable target within the mitochondrial stress pathway and demonstrate that de novo designed proteins can serve as precise molecular tools to dissect DELE1-dependent ISR signaling, with potential for future therapeutic development.

## RESULTS

### Structure-guided design of DELE1 binders that target a critical oligomeric interface

Pharmacological strategies that inhibit the integrated stress response (ISR) have shown therapeutic promise; however, most available inhibitors act on the stress-sensing kinases or the eIF2α translation initiation machinery^13,32–36^. As a result, these approaches broadly suppress ISR signaling across multiple stress inputs rather than selectively modulating the mitochondrial branch. The identification of DELE1 as the mitochondrial stress–specific relay that links mitochondrial dysfunction to HRI activation therefore presents an opportunity to intervene directly at the level of mitochondrial stress signaling. Previous structural studies defined the oligomeric architecture of the DELE1 C-terminal domain (DELE1^CTD^) and demonstrated that DELE1 oligomerization enhances HRI kinase activity and downstream ATF4 induction^22^. Disruption of oligomeric interface I within the first α-helix (α1) of the tetratricopeptide repeat domain prevents DELE1 self-assembly, establishing this helix as a critical regulatory element. We therefore asked whether de novo protein design could be used to generate binders that recognize the α1 helix and block DELE1 oligomerization (**Figure S1A**).

Using the cryo-electron microscopy structure of DELE1^CTD^ (PDB: 8D9X) together with AlphaFold-predicted models as templates, we identified four hydrophobic residues on the α1 helix (F240, L242, F250, and L251) as hotspot positions for binder design (**Figure S1B**). We then used RFdiffusion^37^ to generate backbone scaffolds designed to sterically engage the α1 helix (**Figure 1**). Fifty diffusion steps yielded 83 candidate backbones ranging from 60 to 80 amino acids in length, which were predicted to fold into compact three- or four-helix bundles. Next, ProteinMPNN^38^ was used to design amino acid sequences for each scaffold and to optimize side-chain interactions at the DELE1–binder interface, generating five sequence variants per backbone for a total of 415 designed proteins. To evaluate complex formation and detailed DELE1-binder interactions, we used AlphaFold2 and AlphaFold3 to predict the structures of each DELE1–binder pair^39,40^. Designs were assessed using multiple confidence metrics, including complex root-mean-square deviation (RMSD) relative to the starting model, per-residue confidence score by the predicted Local Distance Difference Test (pLDDT) for the designed protein, and interface pairwise alignment error (PAE_interaction) (**Figure 1 and Table S1**). Based on these metrics, together with manual inspection of predicted complexes, we selected a final set of 12 high-confidence designs for experimental characterization (**Table S1**).

**Figure 1:**
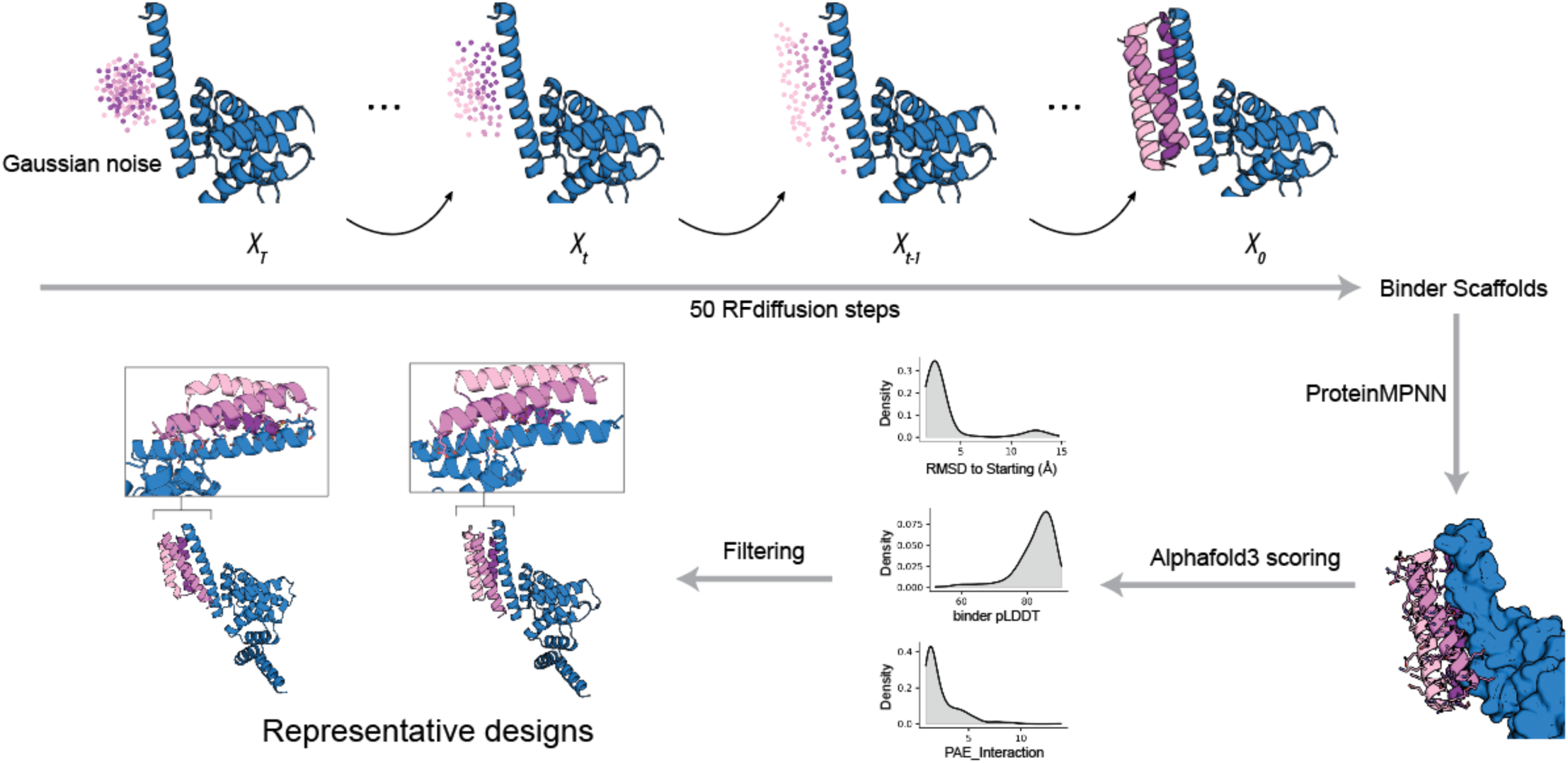
Structure-guided de novo design of DELE1 binders targeting the α1 oligomerization interface. Schematic overview of the de novo protein design pipeline used to generate DELE1 binders. A monomeric DELE1 C-terminal domain (DELE1^CTD^) subunit (red) was used as a template to target the α1 helix that mediates oligomerization in the oligomeric DELE1^CTD^ cryo-EM structure (PDB: 8D9X). RFdiffusion was applied to generate binder backbone scaffolds through iterative denoising steps, starting from Gaussian noise and converging on structured helical scaffolds positioned against the DELE1 α1 helix surface. Candidate backbones were subsequently subjected to sequence optimization using ProteinMPNN. The resulting DELE1–binder complexes were evaluated using AlphaFold2 and AlphaFold3 structure prediction, and designs were filtered based on prediction confidence metrics, including binder RMSD relative to the starting backbone, per-residue confidence (pLDDT), and interface pairwise aligned error (PAE). High-confidence designs were selected for experimental validation.

### Designed binders form stable complexes with DELE1 and disrupt DELE1 oligomerization in vitro

Genes encoding each designed protein were synthesized and cloned into a bacterial expression vector containing an N-terminal 6×His affinity tag. Because the designs are short and frequently lack aromatic residues, a single tyrosine residue was inserted between the 6×His tag and the designed sequence to enable reliable UV detection during chromatographic analysis. To assess whether the designed proteins interact with DELE1 and alter its assembly state, each 6×His-tagged design was co-expressed with maltose-binding protein (MBP)-tagged DELE1 in *E. coli*, followed by nickel affinity purification via the binder tag (**Figure S2A**). SDS–PAGE analysis revealed that MBP-DELE1 co-purified with 11 of the 12 designs, indicating that the majority of designs form stable complexes with DELE1 (**Figure S2B**).

We next examined the oligomeric properties of the 11 co-purified complexes using size-exclusion chromatography (SEC). The majority of DELE1–binder complexes exhibited highly similar SEC profiles, characterized by a dominant, homogeneous peak eluting at approximately 14 mL on a Superdex 200 Increase 10/300 GL column (**Figures 2 and S3**). SDS–PAGE analysis of fractions corresponding to this dominating peak confirmed the presence of both MBP-DELE1 and the designed protein, consistent with formation of a homogenous DELE1–binder complex. One design, binder10, showed reduced co-elution with DELE1 relative to the other binders, indicating weaker complex formation (**Figure S3K, L** and **Table S1**). Nevertheless, for the remaining designs, complex formation was robust and consistent, supporting the conclusion that de novo designed binders can stably associate with DELE1 in vitro. A later-eluting peak at approximately 16 mL contained only the designed protein and likely reflects excess unbound binder that co-purified during nickel affinity purification (**Figures 2 and S3**). To compare the DELE1-binder complexes with oligomeric DELE1 alone, we expressed and purified the MBP-DELE1 in the absence of binder. Consistent with prior observations^22^, DELE1 eluted as a heterogeneous population, with most material appearing in the high–molecular weight region around 10 mL elution volume (**Figures 2 and S3, gray trace**), reflecting its intrinsic propensity to self-assemble into oligomeric species. In contrast, all DELE1–binder complexes eluted at later, lower–molecular weight volumes around 14 mL and were markedly more homogeneous, with noticeably improved yields during purification (**Figures 2 and S3, blue traces**). These results demonstrate that the designed proteins directly associate with DELE1 to form stable complexes and suppress DELE1 oligomerization. This biochemical behavior is consistent with the design model in which the binders engage the α1 helix and prevent formation of the native DELE1 oligomeric scaffold.

**Figure 2:**
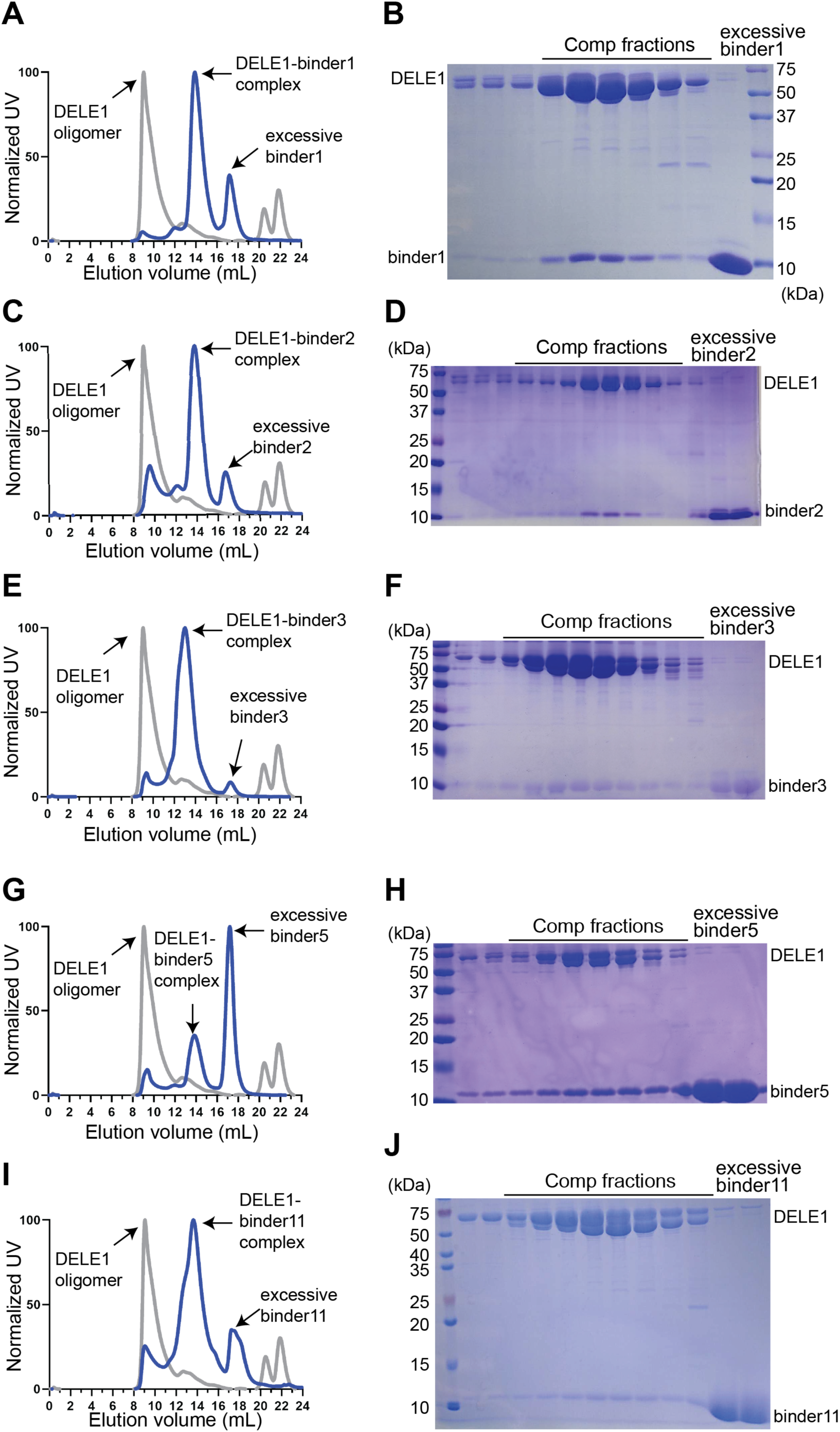
Designed binders directly associate with DELE1 and disrupt its oligomeric state in vitro. (**A, C, E, G, I**) Size-exclusion chromatography (SEC) profiles of MBP-DELE1 co-expressed with individual 6His-tagged binders (binder1, binder2, binder3, binder5, and binder11). Elution profiles show the DELE1–binder complex (blue trace), oligomeric DELE1 species (gray trace), and excess unbound binder (black trace). Elution volumes are indicated on the x-axis. (**B, D, F, H, J**) Coomassie-stained SDS–PAGE analysis of SEC fractions corresponding to the major complex peaks shown in panels A, C, E, G, and I. Fractions from the dominant SEC peak contain both MBP-DELE1 and the corresponding binder, confirming stable complex formation. Lanes labeled “excess binder” contain unbound binder alone. Across all binders shown, co-expression with DELE1 resulted in formation of a homogeneous DELE1–binder complex that eluted at later volumes relative to oligomeric DELE1 alone, consistent with reduced higher-order assembly of DELE1 in vitro.

### Designed binders inhibit mitochondrial stress-induced ISR signaling in cells

We next examined whether the designed proteins that form stable complexes with DELE1 in vitro are capable of modulating ISR signaling in cells. HEK293T cells were transfected with GFP-tagged (all GFP mentioned here and below are mclover) designs, with the GFP fusion enabling detection by immunoblotting because antibodies are not available for the designed sequences. After 24 hours of expression, cells were exposed to either carbonyl cyanide m-chlorophenylhydrazone (CCCP) or oligomycin to induce mitochondrial stress (**Figure 3A**). Both treatments robustly increased ATF4 levels in control cells (**Figure 3B, D**), confirming activation of the DELE1–HRI pathway under these conditions^19,20,41^. Expression of individual designs produced distinct cellular responses. During CCCP treatment, which induces mitochondrial depolarization, 9 of the 11 designs reduced ATF4 accumulation by more than 50% relative to mock-transfected cells, with only binder4 and binder9 exhibiting more modest inhibition (**Figure 3B, C**). During oligomycin treatment, which inhibits mitochondrial ATP synthase, several designs, including binder5, binder11, binder3, binder1, and binder2, also markedly suppressed ATF4 upregulation (**Figure 3D, E**). Together, these data demonstrate that multiple designed proteins, including binder5, binder11, binder3, binder1, and binder2, suppress ISR activation triggered by distinct forms of mitochondrial perturbation.

**Figure 3:**
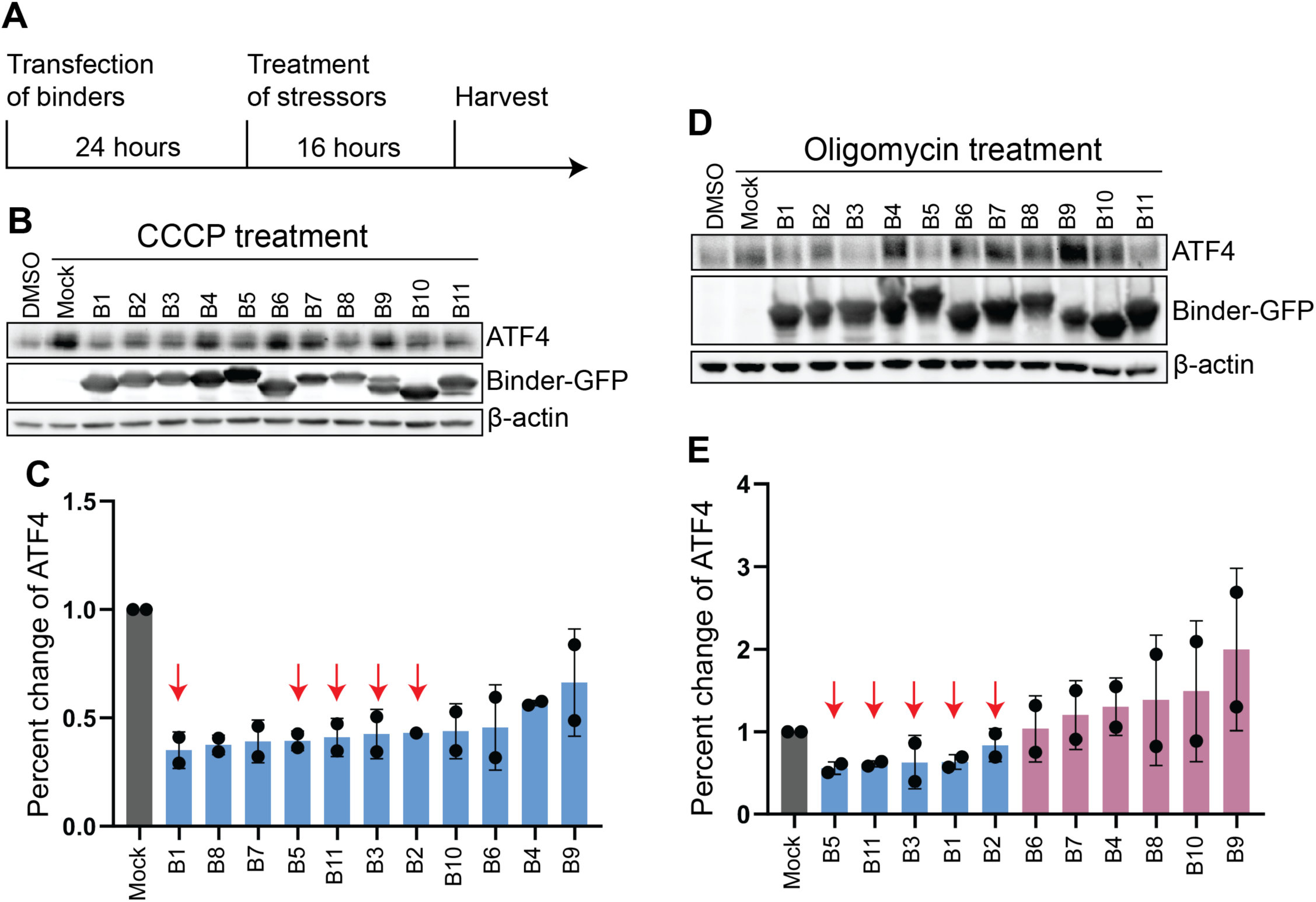
Designed binders inhibit mitochondrial stress-induced ATF4 activation. (**A**) Experimental schematic illustrating the timeline for binder expression and stress treatment. HEK293T cells were transfected with GFP-tagged binders for 24 hours, followed by treatment with mitochondrial stressors for 16 hours prior to harvest. (**B**) Immunoblot analysis of ATF4 levels in HEK293T cells expressing GFP-tagged binders and treated with CCCP. Binder expression was monitored by GFP immunoblotting, and β-actin served as a loading control. **(C)** Quantification of ATF4 levels from CCCP-treated samples shown in panel **B**, normalized to β-actin and expressed relative to mock-transfected cells. Data represent mean ± s.d. from 2 independent experiments. (**D**) Immunoblot analysis of ATF4 levels in HEK293T cells expressing GFP-tagged binders and treated with oligomycin. Binder expression was detected by GFP immunoblotting, with β-actin as a loading control. (**E**) Quantification of ATF4 levels from oligomycin-treated samples shown in panel **D**, normalized to β-actin and expressed relative to mock-transfected cells. Red arrows indicate binders that produced marked suppression of ATF4 induction. DMSO represents cells treated with DMSO and serves as the blank control, rather than treatment with CCCP or oligomycin. Mock represents cells treated with CCCP or oligomycin as a positive control; however, no binders were transfected or overexpressed in these cells. Data represent mean ± s.d. from 2 independent experiments.

To determine whether ISR suppression correlates with direct binder engagement of DELE1, we performed co-immunoprecipitation assays in cells co-expressing FLAG-tagged DELE1 and selected binders. Binder1 and binder5 were chosen because they exhibited the strongest inhibition of ATF4 induction in the CCCP and oligomycin assays, respectively. Both binder1 and binder5 robustly co-precipitated with DELE1, whereas GFP alone did not, confirming that these designs associate with DELE1 in mammalian cells (**Figure S4A**). Notably, binder5 appeared to interact more strongly with DELE1 than binder1, consistent with structural designs indicating that binder5 forms a more extensive hydrophobic interface with DELE1 (**Figure 5C, D**). In addition, DELE1 abundance was increased in whole-cell lysates from binder-expressing conditions compared with GFP controls (**Figure S4A**), suggesting that binder association enhances DELE1 stability or solubility in cells, consistent with our bacterial co-expression results (**Figure 2**). Together, these findings demonstrate that suppression of mitochondrial stress–induced ISR signaling by the designed proteins occurs through their direct physical association with DELE1 in cells.

To evaluate whether ISR inhibition by these designs is specific to mitochondrial stress, we expressed the same subset of binders (binders 1, 2, 3, 5, and 11) in HEK293T cells and treated the cells with thapsigargin, an inducer of endoplasmic reticulum (ER) stress that activates the ISR through the kinase PERK. In contrast to mitochondrial stress conditions, none of the designs affected ATF4 induction following thapsigargin treatment (**Figure S4B, C**). This specificity indicates that the designed proteins selectively interfere with the mitochondrial stress arm of the ISR without perturbing ER stress signaling. Although DELE1 has been established as the essential relay linking mitochondrial stress to HRI activation, its contribution to other ISR branches has remained less investigated. The selective activity of the DELE1-directed designs therefore provides functional evidence that DELE1 operates specifically within the mitochondrial branch of the ISR.

### Designed binders impair adaptive recovery of mitochondrial morphology after transient stress

Mitochondrial morphology remodeling is a central component of the adaptive response to stress, and restoration of elongated tubules following genetic and chemical insults is tightly coupled to the transcriptional and translational programs governed by the ISR^42,43^. Because DELE1 oligomerization is required for efficient activation of HRI and downstream ISR signaling, we asked whether disrupting DELE1-mediated ISR using our designed binders would impair the cell’s ability to re-establish mitochondrial network integrity after transient mitochondrial injury.

Binder1 or binder5 were transiently expressed in U2OS cells, and under basal conditions all samples exhibited the characteristic interconnected mitochondrial network, indicating that expression of the designed proteins alone does not perturb steady-state mitochondrial dynamics (**Figure S5A**). To determine whether binder-mediated inhibition of DELE1 affects the cellular response to acute mitochondrial insult, cells were subjected to a transient CCCP treatment. A 2-hour CCCP exposure induced robust mitochondrial fragmentation in both control and binder-expressing cells (**Figure S5B**), consistent with the well-established sensitivity of mitochondrial morphology to loss of membrane potential^44^. Following washout of CCCP, control cells mounted a rapid adaptive response. Within 4 hours, most cells restored elongated mitochondrial tubules, and by 8 hours the tubular network was largely re-established (**Figure 4A, B**). In marked contrast, expression of either binder1 or binder5 caused a pronounced delay in recovery. At 4 hours post-washout, fewer than 20% of binder-expressing cells reformed elongated mitochondria, representing a substantial deficit relative to controls (**Figure 4A–C**). This impairment persisted at 8 hours, with only approximately 20–25% of binder-expressing cells exhibiting restored networks (**Figure 4A, B, D**), indicating that early steps in the adaptive remodeling process are compromised when DELE1-mediated ISR signaling is inhibited. Notably, mitochondrial morphology eventually recovered by 24 hours across all conditions, consistent with the transient nature of binder expression and the potential existence of DELE1-independent pathways that can support delayed mitochondrial repair.

**Figure 4:**
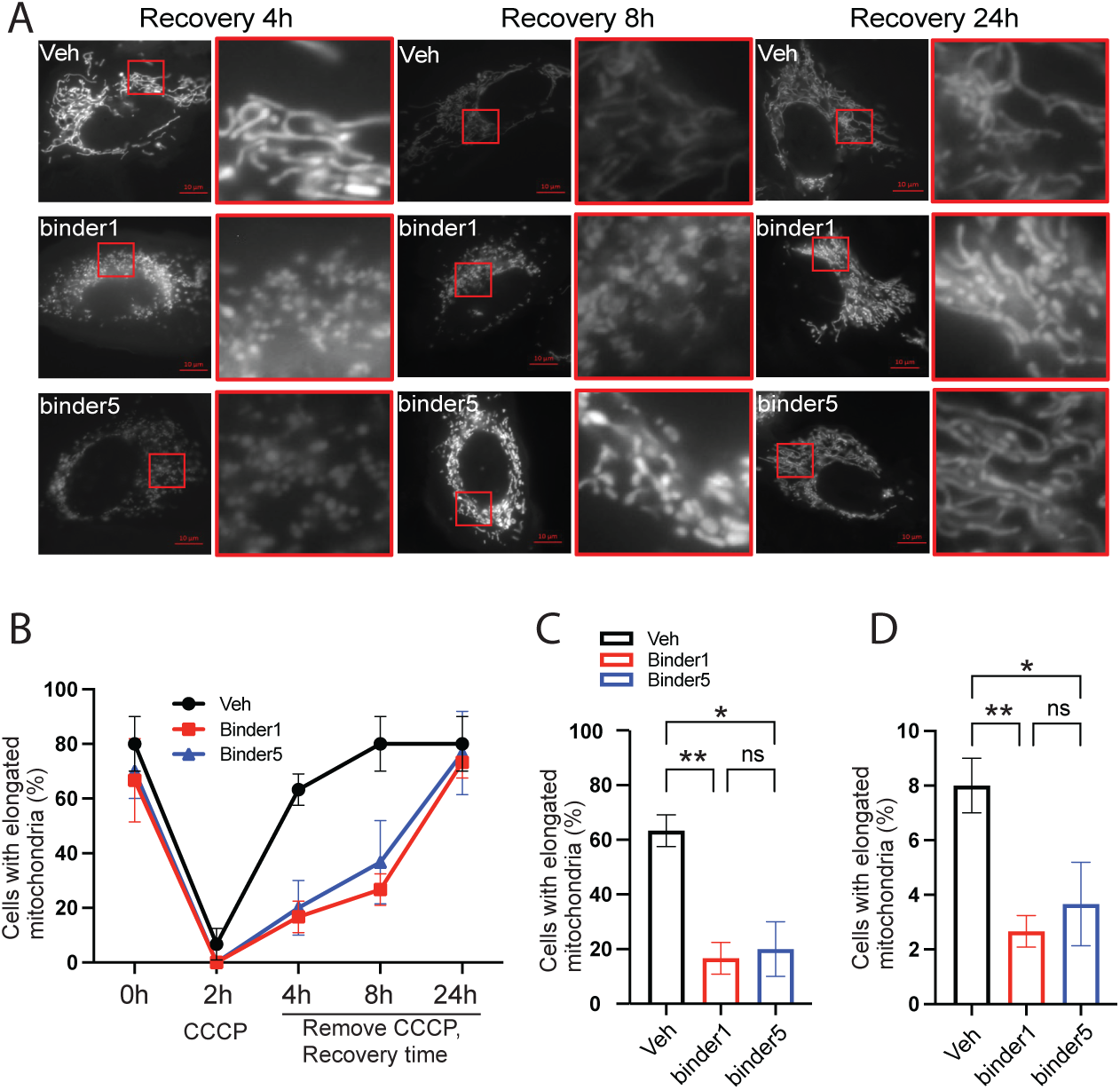
Delayed restoration of mitochondrial network morphology following transient stress in binder-expressing cells. (**A**) Representative fluorescence microscopy images of mitochondrial morphology during recovery following transient CCCP treatment. U2OS cells expressing a mitochondria-targeted pre-CoxIV–BFP reporter alone (Veh) or together with binder1-GFP or binder5-GFP were treated with CCCP for 2 hours, followed by washout and recovery for the indicated times (4h, 8h, and 24h). Insets show 4× magnified views of the regions outlined by red boxes. Scale bars, 20 μm. (**B**) Time course quantification of mitochondrial morphology recovery following CCCP treatment. The percentage of cells exhibiting elongated mitochondrial networks was quantified at baseline (0 h), after 2h CCCP treatment, and at the indicated times (4h, 8h, 24h) following CCCP removal. Control cells (Veh) rapidly restored elongated mitochondrial morphology, whereas cells expressing binder1 or binder5 showed delayed recovery at early time points. Data represent mean ± s.d. from three independent experiments. (**C**) Quantification of the percentage of cells with elongated mitochondria 4h after CCCP removal for the conditions shown in panel A. Bars represent mean ± s.d. from three independent experiments. (**D**) Quantification of the percentage of cells with elongated mitochondria 8 h after CCCP removal for the conditions shown in panel A. Bars represent mean ± s.d. from three independent experiments. Statistical significance was assessed using one-way ANOVA, with significance levels indicated. Veh represents cells treated with as a positive control; however, no binders were transfected or overexpressed in these cells. **: <0.01, *:<0.05, ns: not significant.

The impaired recovery of mitochondrial morphology in binder-expressing cells is consistent with a mechanistic model in which DELE1 oligomerization amplifies mitochondrial stress signals to HRI, and the resulting ISR output facilitates efficient restoration of mitochondrial architecture. By attenuating DELE1-dependent ISR signaling, the designed proteins blunt this adaptive remodeling program, causing mitochondria to remain in a fragmented, injury-associated state for extended periods. Together, these results extend the biochemical and signaling evidence of ISR suppression to a physiological phenotype, demonstrating that DELE1 assembly-state transitions are required not only for ATF4 induction but also for the timely re-establishment of mitochondrial network organization following stress.

### Crystal structure and targeted mutagenesis validate the DELE1-binder interactions

To gain structural insight into how the designed proteins engage DELE1, we attempted to determine the structure of the DELE1–binder complex. Crystallization of the complex was unsuccessful, likely due to conformational flexibility introduced by the MBP fusion required to stabilize DELE1. Cryo-EM was also not feasible because the complex falls below the practical size limit for high-resolution single-particle analysis, and the flexible MBP tag further introduced substantial technical challenges for particle alignment of an already small complex of approximately 40 kDa. We therefore focused on the designed proteins themselves and successfully determined the crystal structure of binder5 at 2.6 Å resolution (**Figure 5A**). The structure revealed a well-packed four-helix bundle that closely matched the design model, with an RMSD of 1.5 Å relative to the computational model, providing direct experimental validation of the accuracy of the design pipeline (**Figure 5B**). Notably, the observed structural differences were primarily confined to side-chain conformations rather than backbone rearrangements (**Figure S6A**), indicating that the overall fold and topology were accurately specified during design.

**Figure 5:**
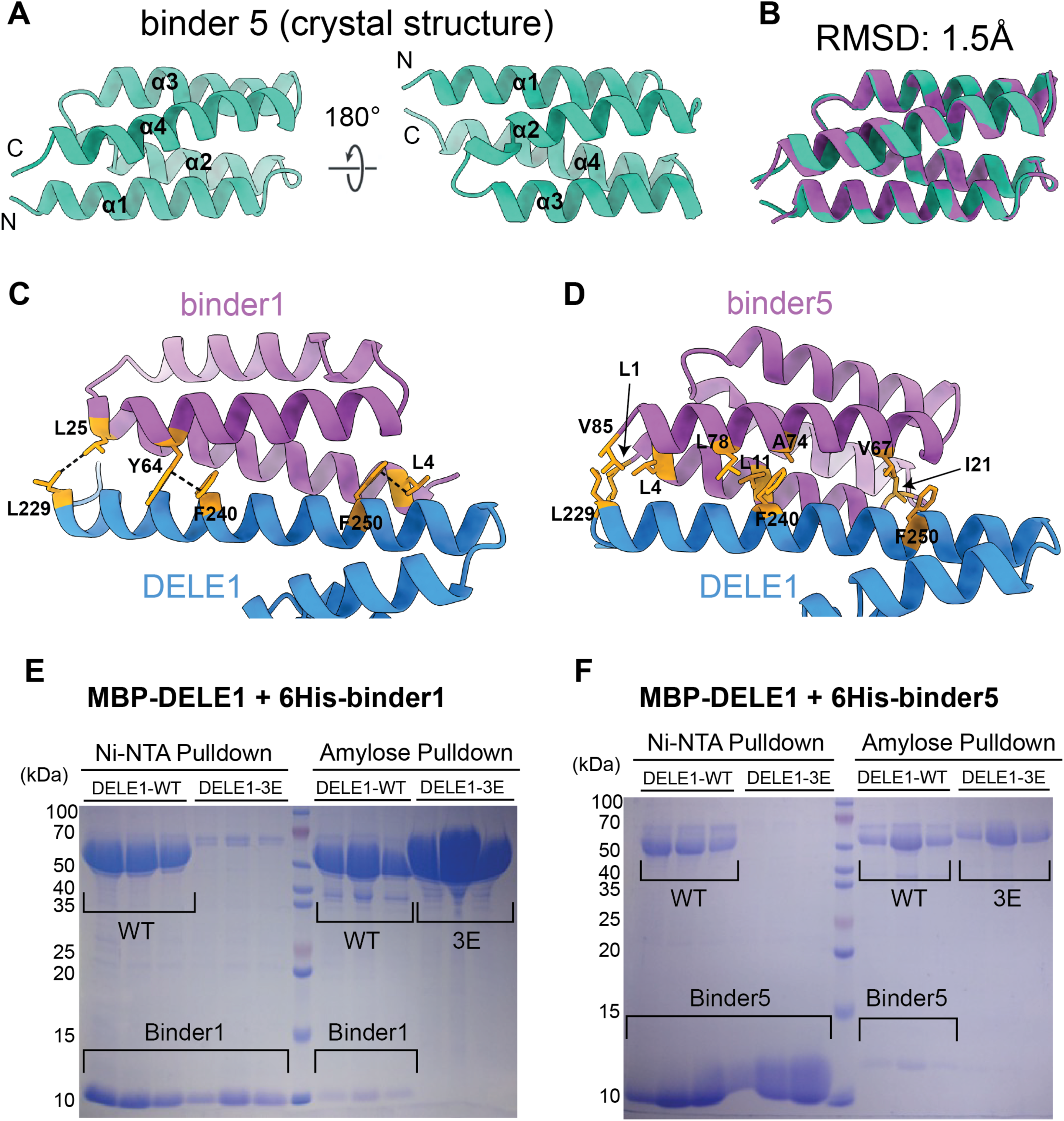
Structural and mutational validation of the DELE1–binder interaction interface. (**A**) Crystal structure of binder5 determined at 2.6 Å resolution, shown in two orientations related by a 180° rotation. The structure adopts a compact four-helix bundle fold, with N- and C-termini indicated. (**B**) Superposition of the binder5 crystal structure (teal) with the design model (purple), showing close agreement between the experimental structure and the predicted fold (RMSD = 1.5 Å). (**C**) Designed model of binder1 (purple) bound to the DELE1 α1 helix (blue). Side chains of key hydrophobic residues on DELE1 (L229, F240, and F250) and interacting residues on binder1 are shown and labeled. Dashed lines indicate predicted side-chain interactions. **(D)** Designed model of binder5 (purple) bound to the DELE1 α1 helix (blue). Residues forming the predicted hydrophobic interaction interface on both DELE1 and binder5 are shown and labeled. (**E**) Affinity pulldown analysis of MBP-DELE1 (wild-type or L229E/F240E/F250E triple mutant, “3E”) co-expressed with 6His-tagged binder1. Ni–NTA pulldown via the binder tag (left) and reciprocal amylose pulldown via MBP-DELE1 (right) show robust co-purification with wild-type DELE1 but loss of interaction with the 3E mutant. (**F**) Affinity pulldown analysis of MBP-DELE1 (wild-type or 3E) co-expressed with 6His-tagged binder5, performed as in panel E. Binder5 co-purifies with wild-type DELE1 but not with the 3E mutant.

Structural models of binder1 and binder5 in complex with DELE1 revealed that both designs engage the same hydrophobic face along the DELE1 α1 helix. In the binder1–DELE1 model (**Figure 5C, Figure S1C, D**), DELE1 residues L229, F240, and F250 form a contiguous hydrophobic patch that packs against binder1 residues L25, Y64, and L4. A prominent feature of this interface is a π–π stacking interaction between the aromatic ring of binder1-Y64 and DELE1-F240, adopting a face–edge geometry that stabilizes the central region of the complex. At the N-terminal end of the α1 helix, binder1-L25 packs against DELE1-L229, whereas binder1-L4 engages DELE1-F250 toward the C-terminal end, collectively forming a three-point hydrophobic clamp that spans the length of the α-helical surface. The binder5–DELE1 model (**Figure 5D**) revealed an even more extensive apolar interface. Binder5 residues V85, L1, L4, L78, L11, A74, V67, and I21 form a continuous hydrophobic ridge that aligns along the L229–F240–F250 axis of DELE1. Binder5-V85, L1 and L4 pack against DELE1-L229, while binder5-L78, L11, and A74 contour around DELE1-F240, producing broad aromatic–aliphatic complementarity despite the absence of a π–π stacking interaction. Toward the C-terminal region of the α1 helix, binder5-V67, and I21 engages DELE1-F250 in a tightly packed hydrophobic interaction. Although binder5 lacks an aromatic residue positioned for π–π stacking, the greater number and continuity of hydrophobic contacts across the α1 surface result in a more expansive interaction footprint than that of binder1, consistent with the stronger binding observed in cellular co-immunoprecipitation assays.

To experimentally test these structural models, we generated a DELE1 triple mutant in which the predicted binder-interacting residues were substituted with glutamate (L229E, F240E, and F250E; hereafter referred to as DELE1-3E) to disrupt the hydrophobic interaction surface while preserving the overall helical character of α1. MBP-tagged DELE1 (WT or 3E) was co-expressed with 6His-tagged binder1 or binder5 in *E. coli*, and complex formation was assessed using two reciprocal affinity pulldown assays. In Ni–NTA pulldowns, wild-type DELE1 robustly co-eluted with 6×His-binder1, whereas the DELE1-3E mutant failed to co-purify, indicating a complete loss of interaction (**Figure 5E, F**). Reciprocal MBP pulldowns yielded the same result, with binder1 enriched only in the presence of wild-type DELE1. Binder5 behaved analogously, showing strong co-elution with wild-type DELE1 but no detectable association with the DELE1-3E mutant in either pulldown assay (**Figure 5E, F**). Importantly, the DELE1-3E mutant expressed well and eluted at a smaller apparent molecular weight than wild-type DELE1 during SEC (**Figure S6B–D**), consistent with prior reports that residues within this hydrophobic triad contribute to DELE1 oligomerization rather than global protein stability. The absence of binder association in the 3E mutant background therefore reflects a specific disruption of the designed interaction interface rather than misfolding or reduced stability of DELE1. Together, these biochemical and structural results validate the computational models, establishing L229, F240, and F250 as essential DELE1 contact residues for both binder1 and binder5 and confirming the hydrophobic interaction network that underlies the de novo design strategy.

### The DELE1-HRI association is not disrupted by the designed binders

Previous studies demonstrated that DELE1 associates with the N-terminal domain of HRI (HRI^NTD^: residues 1–160) in cells using co-immunoprecipitation^22^, but whether this interaction is direct and whether it is perturbed by the designed proteins remained unclear. To address these questions, we purified HRI^NTD^ and analyzed its interactions with DELE1 and DELE1–binder complexes using SEC. Initial attempts to express HRI^NTD^ alone in bacteria yielded highly insoluble protein; however, fusion to an N-terminal MBP-TEV tag restored solubility and enabled purification to homogeneity (**Figure S7A, B**). Surprisingly, MBP–TEV-HRI^NTD^ eluted as a homogeneous but early-eluting, high–molecular weight species, suggesting that the isolated HRI N-terminal domain may form higher-order assemblies in solution (**Figure S7A, B**). The oligomeric properties of HRI^NTD^ and their potential relevance to ISR regulation remain to be investigated. By contrast, MBP–DELE1–binder1 and MBP–DELE1–binder5 complexes eluted at approximately 14 mL, consistent with their behavior in earlier SEC analyses (**Figures 6A, C, black traces; Figure 2**).

**Figure 6:**
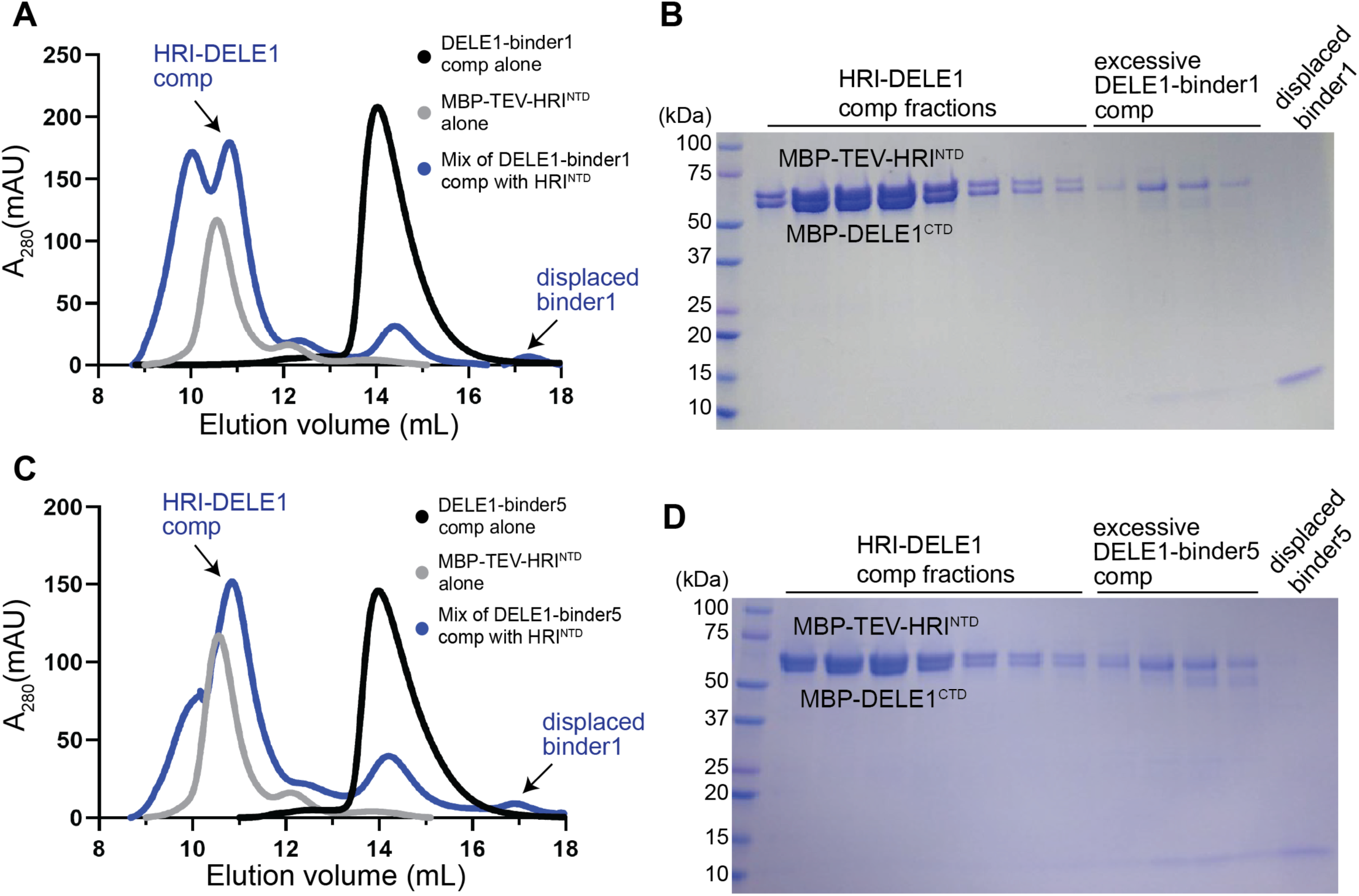
Designed binders do not disrupt the DELE1-HRI interaction. (**A**) SEC analysis of MBP–TEV–HRI^NTD^ alone (gray trace), the purified DELE1–binder1 complex (black trace), and the mixture of DELE1–binder1 with MBP–TEV–HRI^NTD^ (blue trace). Upon mixing, the DELE1–binder1 peak is reduced and replaced by earlier-eluting high–molecular weight species corresponding to DELE1–HRI complexes, while free binder1 appears as a later-eluting peak. (**B**) Coomassie-stained SDS–PAGE analysis of SEC fractions from the mixture (blue trace) in panel A. Fractions corresponding to the early-eluting peaks contain both MBP–DELE1-CTD and MBP–TEV-HRI^NTD^, confirming formation of DELE1–HRI complexes. Binder1 is absent from these fractions and detected only in later fractions corresponding to displaced binder. (**C**) SEC analysis performed as in panel A using the DELE1–binder5 complex. Mixing with MBP–TEV–HRI^NTD^ similarly yields early-eluting DELE1–HRI species and displacement of binder5. (**D**) SDS–PAGE analysis of SEC fractions from the mixture (blue trace) in panel C confirming the formation of DELE1-HRI complexes and the displaced binder5.

When MBP-TEV–HRI^NTD^ was mixed with the purified DELE1–binder1 complex and subjected to SEC, the resulting chromatogram exhibited two distinct high–molecular weight species accompanied by a reduction in the original DELE1–binder1 peak (**Figure 6A, blue trace**). SDS–PAGE analysis of the corresponding fractions revealed that both high–molecular weight species contained MBP-DELE1^CTD^ and MBP–HRI^NTD^, demonstrating that HRI^NTD^ forms a stable complex with DELE1 in vitro (**Figure 6B**). To further validate formation of the DELE1–HRI complex, fractions corresponding to the high–molecular weight species were treated with TEV protease. Because the MBP-TEV-HRI^NTD^ construct contains a TEV cleavage site between the MBP tag and HRI^NTD^, TEV treatment selectively removed the MBP tag from HRI-NTD, but not from the non-cleavable MBP-DELE1^CTD^. Following cleavage, the liberated HRI^NTD^ (∼20 kDa) remained associated with MBP-DELE1^CTD^ and continued to elute in the early SEC fractions (**Figure S7C, D**), providing additional confirmation of a direct DELE1–HRI interaction. In contrast, binder1 was completely absent from these early-eluting fractions and instead appeared exclusively in the low–molecular weight region (**Figure 6A, B**), indicating full displacement of binder1 from DELE1 upon HRI binding. To further confirm that binder1 does not interfere with DELE1–HRI association, excess binder1 was added to a preformed DELE1–HRI complex and the mixture was analyzed by SEC followed by SDS–PAGE (**Figure S7E, F**). Under these conditions, excess binder1 did not disrupt the DELE1–HRI complex, suggesting that binder1 does not substantially occlude the HRI binding interface.

A similar outcome was observed for binder5. Mixing MBP-TEV-HRI^NTD^ with the DELE1–binder5 complex yielded high–molecular weight SEC species containing both DELE1 and HRI^NTD^, whereas binder5 eluted exclusively in the low–molecular weight region, indicating complete displacement (**Figure 6C, D**). Addition of excess binder5 to a preformed DELE1–HRI complex likewise failed to disrupt complex formation, as assessed by SEC and SDS–PAGE (**Figure S7G, H**), confirming that binder5 does not interfere with DELE1–HRI association. We consistently observed the appearance of two DELE1–HRI species likely reflects distinct assembly states or stoichiometries of the DELE1–HRI complex formed in vitro (**Figure 6C, blue trace**). Together, these results demonstrate that HRI^NTD^ binds directly to DELE1 and that this interaction is not occluded or destabilized by the designed proteins. Instead, the designed proteins inhibit ISR signaling by blocking DELE1 oligomerization while preserving DELE1–HRI recognition and binding.

## DISCUSSION

Mitochondrial stress signaling through the DELE1–HRI axis is a fundamental surveillance pathway that safeguards cellular proteostasis and adapts translational output to organellar dysfunction. Although DELE1 has been firmly established as the dedicated relay that transmits mitochondrial stress signals to the cytosolic ISR machinery, the field has lacked molecular tools capable of selectively modulating DELE1 function without perturbing other ISR branches. Here, we address this critical gap by generating de novo designed protein binders that specifically target the α1 helix of DELE1, a structural element required for oligomerization and signal propagation. Through an integrated combination of computational design, X-ray crystallography and structural modeling, biochemical analyses, and cell-based functional assays, we demonstrate that these binders disrupt DELE1 assembly, attenuate mitochondrial stress–induced ISR signaling, and impair downstream recovery of mitochondrial morphology. These findings establish DELE1 as a druggable signaling target and highlight the power of rational protein design to precisely manipulate ISR pathway components.

Our binders also function as highly specific tools for dissecting mitochondrial stress signaling. In contrast to kinase inhibitors or genetic ablation, which globally alter ISR activity and confound pathway-specific interpretation, the designed proteins target a single structural element of DELE1: the α1 oligomerization interface. By selectively disrupting DELE1 assembly while preserving its mitochondrial targeting, OMA1-dependent cleavage, cytosolic accumulation, and HRI engagement, the binders achieve a level of pathway specificity that has not been previously accessible. This enables precise interrogation of how DELE1 assembly-state transitions encode stress signals and amplify HRI activation, providing a mechanistic framework for understanding how mitochondrial perturbations are transmitted to the translational machinery. This level of target precision opens new opportunities for interrogating DELE1 function in normal physiology, mitochondrial pathology, and tissue-specific stress adaptation. More broadly, our design framework provides a blueprint for engineering selective modulators of signaling proteins whose activation depends on conformational or oligomeric switching.

At the molecular level, our study defines a hydrophobic triad (L229, F240, and F250) along the DELE1 α1 helix as the core anchoring surface for binder engagement. The binder5 crystal structure, combined with AlphaFold3 modeling and targeted mutagenesis, confirms that this triad constitutes a discrete and druggable interface. By validating that disruption of these residues abolishes binder binding and alters DELE1 assembly, we establish a mechanistic foundation for understanding and how DELE1 self-association contributes to ISR activation. These insights complement recent structural studies and begin to outline an emerging picture in which mitochondrial stress induces a controlled reorganization of DELE1 architecture that in turn modulates HRI activation.

Our findings also refine the mechanistic relationship between DELE1 oligomerization and HRI recruitment. Although the binders potently block DELE1 assembly, they are fully displaced by HRI-NTD during competition experiments using purified proteins. These results suggest that HRI engages DELE1 with substantially higher affinity than the binders and suggests that the DELE1–HRI interface may either partially overlap the α1 surface or trigger an allosteric rearrangement that reduces binder affinity. We propose a hierarchical model in which DELE1 oligomerization generates a competent docking platform for HRI, and HRI binding drives the transition toward a fully active signaling complex. The presence of two high–molecular weight DELE1–HRI assemblies suggests that DELE1 and HRI may traverse multiple intermediate states prior to kinase activation, a process that remains to be structurally and mechanistically defined.

The biochemical behavior of HRI-NTD also raises intriguing biological implications. MBP–HRI-NTD elutes as a high–molecular weight species, hinting that the isolated N-terminal region may possess an intrinsic propensity for self-association. In cellular contexts, HRI activation requires dimerization and trans-autophosphorylation of its C-terminal kinase domain. The higher-order state of the HRI-NTD observed in vitro may therefore reflect an inherent clustering mechanism that facilitates rapid activation upon DELE1 engagement. Whether DELE1 recruits pre-assembled HRI units, stabilizes weak HRI clustering, or directly organizes an activation-competent HRI superstructure remains an open and compelling question.

Finally, our work provides a platform for future exploration of DELE1 biology and therapeutic modulation of the mitochondrial ISR. Determining the high-resolution structures of the DELE1–binder and DELE1–HRI complexes will be essential for deciphering how assembly-state transitions couple stress signals. Optimization of binder affinity, stability, and intracellular delivery may enable next-generation protein therapeutics aimed at diseases driven by chronic mitochondrial stress and maladaptive ISR activation. Taken together, our study delivers conceptual, structural, and mechanistic advances in mitochondrial stress signaling and demonstrates the feasibility of using de novo protein design to engineer selective modulators of intracellular surveillance pathways.

### limitations of the study

Although this work establishes de novo designed binders as powerful and highly specific modulators of DELE1, several limitations remain. First, we were unable to determine the high-resolution structure of the DELE1–binder complex due to challenges in crystallizing MBP-tagged DELE1 and the small size of the complex for cryo-EM, which limits direct visualization of binder engagement and its structural consequences on DELE1. Second, while our biochemical and cellular assays demonstrate that the binders disrupt DELE1 oligomerization and attenuate mitochondrial stress–induced ISR activation, their affinities, stoichiometries, and dynamic behavior in living cells remain to be fully characterized. Third, although our binders serve as valuable research tools, further engineering will be required to optimize stability, delivery, and potency for potential therapeutic applications. Finally, all functional studies were performed in cultured cells, and it will be important to evaluate the physiological relevance and therapeutic potential of DELE1 binders in vivo, particularly in disease models characterized by chronic mitochondrial stress or maladaptive ISR activation. These limitations outline key directions for future work but do not alter the conclusions of this study.

## DATA DEPOSITION

The X-ray crystallographic coordinates and structure factors have been deposited in the Protein Data Bank (PDB) under accession code 9ZMT. The predicted structural models of the DELE1-binder1 and DELE1-binder5 complexes have been deposited into ModelArchive (https://modelarchive.org) with accession ID ma-31vqf and ma-ytheg, respectively.

## ACKNOWLEDGMENTS

We thank the members of the Yang Lab for insightful discussions and comments throughout this work. X-ray diffraction data were collected at the Southeast Regional Collaborative Access Team (SER-CAT) 22-ID-D beamline at the Advanced Photon Source, Argonne National Laboratory. SER-CAT is supported by its member institutions, equipment grants S10_RR25528, S10_RR028976, and S10_OD027000 from the National Institutes of Health, and by funding from the Georgia Research Alliance. We thank Dr. Shangming Tang at the University of Virginia for sharing the mammalian cell lines. We thank Dr. Tsan Sam Xiao at Case Western Reserve University for kindly sharing the MBP tagged vectors. This work was supported by a pilot grant from the American Cancer Society and Cancer Center Research Gift Funding through the University of Virginia Comprehensive Cancer Center (Grant #134088-IRG-19-143-33-IRG) awarded to J.Y.. B.K. is supported by NIH grant R35GM131923. D.F.K. is supported by NCI grant 1U54CA274499.

## AUTHOR CONTRIBUTIONS

R.Y. and D.Y. performed the protein biochemical studies with assistance from K.P.L. R.Y. determined the X-ray crystal structure. K.Z. and Y.W. conducted the cell-based functional studies with assistance from A.A.P. M.M. and B.K. performed the computational protein design. D.F.K. provided reagents for the mitochondrial morphology assays. J.Y. conceived and supervised the project, interpreted the data, and wrote the manuscript. R.Y., K.Z. and Y.W. assisted in preparing the figures. All authors reviewed and commented on the manuscript.

## METHODS

### Reagents, plasmids and cell lines

Genes encoding the designed protein sequences were synthesized by Twist Biosciences (South San Francisco, CA). Each design was cloned into a modified pET28a bacterial expression vector containing an N-terminal 6×His affinity tag. A single tyrosine residue was engineered between the 6×His tag and the designed protein sequence. The C-terminal domain of human DELE1 (DELE1^CTD^, residues K225–G515) was cloned into a modified pET20b bacterial expression vector containing a non-cleavable N-terminal MBP tag and a C-terminal 6×His tag. All cloning was performed using Gibson Assembly. For co-expression experiments, DELE1^CTD^ and the DELE1-3E mutant were cloned into a similar vector in which the C-terminal 6×His tag was removed using Q5 site-directed mutagenesis (New England Biolabs, #E0554S). Point mutations L229E, F240E, and F250E were introduced using Q5 mutagenesis with the wild-type DELE1^CTD^ construct as the template. The N-terminal domain of human HRI (residues 2–160) was cloned into a modified pET20b vector containing an N-terminal 8×His tag and an MBP tag that is cleavable by TEV protease. TEV protease was expressed and purified in house. HEK293T and U2OS cells were obtained as gifts from Dr. Shangming Tang’s laboratory and were cultured in Dulbecco’s modified Eagle medium (DMEM, Gibco, #11965-092) supplemented with 10 percent fetal bovine serum (Corning, #35-015-CV) and penicillin–streptomycin (Gibco, #15140-122). Cells were passaged using 1× DPBS (Gibco, #14190-144) and 0.05 percent trypsin–EDTA (Gibco, #25300-054).

### Computational de novo design of DELE1 binders

De novo protein binders were designed to target the α1 helix of the human DELE1 C-terminal domain (DELE1^CTD^), which mediates oligomerization in the activated DELE1 assembly. A monomeric DELE1^CTD^ subunit was used as the design target, with the α1 helix explicitly defined as the binding “epitope”. Binder backbone scaffolds were generated using RFdiffusion. The DELE1 α1 helix surface was provided as a fixed conditioning input, and binder backbones were generated through iterative denoising steps starting from Gaussian noise. During diffusion, structural constraints guided the emergence of compact, predominantly helical binder scaffolds positioned against the α1 interface with appropriate surface complementarity and orientation. This process produced diverse backbone geometries converging on the DELE1 α1 helix surface. Candidate binder backbones generated by RFdiffusion were subjected to sequence design and optimization using ProteinMPNN. Amino acid sequences were designed to stabilize the binder fold while optimizing interactions at the DELE1–binder interface. Multiple sequence variants were generated for each backbone to explore sequence space while preserving the designed binding geometry. Designed DELE1–binder complexes were evaluated using AlphaFold2 and AlphaFold3 structure prediction. For each design, predicted complex structures were analyzed to assess reproducibility of the intended binding mode and interface integrity. Designs were filtered based on prediction confidence metrics, including binder backbone RMSD relative to the RFdiffusion-generated scaffold, per-residue confidence scores (pLDDT), and PAE. Designs exhibiting large structural deviations, low confidence at the interface, or poorly defined DELE1–binder contacts were excluded. High-confidence designs that consistently reproduced the intended α1 helix binding mode across AlphaFold predictions were selected for experimental validation. A final set of 12 de novo designs were advanced for biochemical, structural, and cellular characterization.

### Protein expression and purification

*Escherichia coli* Rosetta (DE3) cells (Agilent Technologies, #70954) were used for all recombinant protein expression. Plasmids encoding wild-type MBP–DELE1^CTD^, MBP–TEV–HRI^NTD^, or individual de novo designed binders were transformed into Rosetta (DE3) cells. Cultures were grown in Luria Broth (Miller) medium at 37 °C until the optical density at 600 nm reached ∼0.8. Protein expression was induced by the addition of 1 mM IPTG, and cultures were incubated at 16 °C overnight. Cells were harvested by centrifugation at 4,000g for 20 min at 4 °C and resuspended in lysis buffer containing 25 mM Tris-HCl (pH 8.0), 500 mM NaCl, 10% glycerol, and 100 µM PMSF. PMSF was omitted during purification of the designed binders due to their stability in Tris-HCl buffer. Cells were lysed by sonication, and lysates were clarified by centrifugation at 25,000g for 30 min at 4 °C. Cleared lysates were subjected to Ni–NTA affinity chromatography using Ni–NTA resin (Cytiva, #17531803) equilibrated in lysis buffer and loaded onto gravity flow columns. After sample loading, columns were washed sequentially with 10 column volumes of wash buffer A, consisting of lysis buffer supplemented with 25 mM imidazole, followed by 5 column volumes of wash buffer B containing lysis buffer supplemented with 50 mM imidazole. Bound proteins were eluted with elution buffer containing 25 mM Tris-HCl (pH 8.0), 500 mM NaCl, 10% glycerol, and 300 mM imidazole. For co-expression experiments used in initial binding tests, plasmids encoding MBP–DELE1^CTD^ and individual de novo designed binders were co-transformed into E. coli cells. Expression conditions were identical to those used for single-protein expression. During subsequent Ni–NTA purification, all buffers contained 150 mM NaCl instead of 500 mM NaCl. MBP–DELE1-3E alone was expressed under the same conditions as other proteins. Purification was performed using amylose affinity chromatography with amylose resin (Cytiva, #17531803) using a gravity flow setup. After binding, the resin was washed with 10 column volumes of lysis buffer and proteins were eluted with lysis buffer supplemented with 30 mM maltose. All proteins were further purified by size-exclusion chromatography using a Superdex 200 Increase 10/300 GL column (Cytiva) on an ÄKTA pure protein purification system (Cytiva). The size-exclusion chromatography buffer consisted of 25 mM Tris-HCl (pH 8.0), 150 mM NaCl, and 5% glycerol. Peak fractions corresponding to target proteins were pooled, concentrated, aliquoted, flash frozen in liquid nitrogen, and stored at −80 °C.

### Crystallization, data collection, and structure determination

The excessive binder5 from the co-expression and purification of the DELE1-binder5 complex was used for crystallization. The binder5 protein was in the buffer containing 25 mM Tris-HCl (pH 8.0), 150 mM NaCl, and 5% glycerol. Crystallization screening was set up using the vapor-diffusion method (sitting drop) on the Intelli-Plate 96-3 low-profile crystallization plate (Art Robbins Instruments, #102-0001-13) by mixing the protein at 11.7 mg/mL with an equal volume of reservoir solution. The best crystals sent for data collection were grown with a solution containing 0.1 M Bis-Tris pH 6.0, 0.2 M Ammonium Acetate, 45% v/v (+/-)-2-Methyl-2,4-pentanediol (MPD) at 4°C. The needle to rod-shape crystal came into formation in 5-7 days. All crystals were flash frozen in liquid nitrogen with the reservoir solution with 45% MPD as cryoprotectant. Diffraction datasets were collected on 22-ID-D beamlines (SER-CAT) at Advance Photon Source, Argonne National Laboratory. Three datasets of binder5 were collected from different spots on one slender crystal and were individually indexed and integrated with DIALS^44^ and then scaled and merged with the CCP4i2^45^. The crystal belongs to space group C2221 (a = 44.786 Å, b = 114.889 Å, c = 79.214 Å, α = β = γ = 90°) The structure was determined by molecular replacement using a predicted model generated by the AlphaFold 3.0 server. Model building was performed in COOT^46^ and further improvement was carried out with rounds of refinement using Phenix.refine^47^ and model tuning via COOT. The refined model corresponds to the highest resolution of 2.6 Å. The main chain fits well with the electron density and plenty of side chains are visible in the map. The data collection and refinement statistics for this structure are listed in Table 1.

**Table 1:** Crystallography data collection and refinement statistics. Values in parentheses are data for the highest resolution shell.

|  |  |
| --- | --- |
| Macromolecule | DELE1 binder 5 (PDB ID 9ZMT) |
| Wavelength (Å) | 1 |
| Resolution range (Å) | 57.44 - 2.6 (3.28 - 2.6) |
| Space group | C 2 2 21 |
| Unit cell (a, b, c, Å) | 44.786 114.889 79.214 90 90 90 |
| Total reflections | 251285 (123725) |
| Unique reflections | 6585 (3229) |
| Multiplicity | 38.2 (38.3) |
| Completeness (%) | 99.77 (99.72) |
| Mean I/sigma(I) | 11.36 (5.36) |
| Wilson B-factor | 42.17 |
| R-merge | 0.4741 (1.435) |
| R-meas | 0.4808 (1.456) |
| R-pim | 0.07842 (0.2374) |
| CC1/2 | 0.961 (0.527) |
| CC* | 0.99 (0.831) |
| Reflections used in refinement | 6571 (3220) |
| Reflections used for R-free | 334 (175) |
| R-work | 0.4517 (0.4447) |
| R-free | 0.4833 (0.4768) |
| Number of non-hydrogen atoms | 682 |
| macromolecules | 682 |
| ligands | 0 |
| solvent | 0 |
| Protein residues | 85 |
| RMS(bonds) | 0.013 |
| RMS(angles) | 2.42 |
| Ramachandran favored (%) | 90.36 |
| Ramachandran allowed (%) | 9.64 |
| Ramachandran outliers (%) | 0 |
| Rotamer outliers (%) | 9.86 |
| Clashscore | 31.01 |
| Average B-factor | 49.6 |
| macromolecules | 49.6 |
| Number of TLS groups | 1 |

### Ni-NTA and amylose resin pulldown assay

Ni–NTA and amylose resin pulldown assays were performed as part of the structure-guided mutagenesis experiments. Wild-type DELE1^CTD^ and the DELE1^CTD^(3E) mutant were cloned into a modified pET20b vector containing a non-cleavable N-terminal MBP tag. Plasmids encoding MBP–DELE1^CTD^ (wild type or 3E mutant) were co-transformed with plasmids encoding Binder1 or Binder5 into Escherichia coli Rosetta (DE3) cells. Bacterial culture, induction, and lysis were performed as described above. Following cell lysis and clarification, supernatants from each co-expression were adjusted to equal volumes using lysis buffer and divided equally into two halves. One half was applied to 0.5 mL bed volume of Ni–NTA resin, and the other fraction was applied to 0.5 mL bed volume of amylose resin. Binding, washing, and elution steps were performed using the same buffer compositions and conditions described for affinity purification. Elution fractions of 0.5 mL were collected, and the first three elution fractions from each pulldown experiment were analyzed by SDS–PAGE.

### Complex formation and displacement assay

MBP–TEV–HRI^NTD^ and MBP–DELE1^CTD^:binder1 or MBP–DELE1^CTD^:binder5 complexes were purified separately as described above. Purified MBP–TEV–HRI^NTD^ was mixed with the MBP–DELE1^CTD^:binder1 or MBP–DELE1^CTD^:binder5 complex at a 1:1 molar ratio. Protein mixtures were incubated on ice for 30 min, diluted to a final volume of 500 µL with size-exclusion chromatography buffer, and injected onto an ÄKTA pure chromatography system for size-exclusion chromatography using a Superdex 200 10/300 GL column. Peak fractions from the size-exclusion chromatography runs of the mixed samples were collected and analyzed by SDS–PAGE. As a negative control, MBP–HRI^NTD^ at the same molar amount used in the mixing experiments was incubated on ice for 30 min and analyzed by size-exclusion chromatography and SDS–PAGE under identical conditions. Formation of the HRI^NTD^:DELE1^CTD^ complex was further validated by TEV protease cleavage. To test whether binder1 or binder5 can disrupt the DELE1-HRI complex, The newly formed MBP–TEV–HRI^NTD^:MBP–DELE1^CTD^ complex were pooled, concentrated, and mixed with purified binder1 or binder5 at a molar ratio of 1:4. These samples containing excess binder were incubated on ice for 30 min and subsequently analyzed by size-exclusion chromatography and SDS–PAGE.

### Transient transfection

HEK293T cells were seeded in six-well plates at a density of 2.5 × 10^5^ cells per well in 2 mL of complete Dulbecco’s modified Eagle medium and cultured for 48 h to reach approximately 50–70 percent confluence on the day of transfection. For each well, 2.5 µg of binder–GFP plasmid DNA was diluted in 125 µL of Opti-MEM I reduced-serum medium (Thermo Fisher Scientific, #31985-070) and mixed with 5 µL of P3000 reagent (Thermo Fisher Scientific, L3000008) by gentle pipetting. In a separate tube, 5 µL of Lipofectamine 3000 reagent was diluted in 125 µL of Opti-MEM I. The diluted Lipofectamine 3000 reagent was added to the DNA–P3000 mixture, gently mixed, and incubated for 15 min at room temperature. Following incubation, the transfection mixture, with a total volume of 250 µL per well, was added dropwise to the cells, which were gently rocked to ensure even distribution. Cells were returned to a 37 °C incubator with 5 percent CO₂ and maintained for 24 h prior to downstream drug treatment assays.

### Drug treatments

HEK293T cells were treated 24 h after transient transfection. For each well of a six-well plate, drug solutions were prepared by adding either 4 µL of 10 mM CCCP, 4 µL of oligomycin stock solution to 2 mL of fresh Dulbecco’s modified Eagle medium, followed by thorough mixing. For thapsigargin treatment, drug solutions were prepared by adding 1.2µL stock solution to 1 mL of fresh Dulbecco’s modified Eagle medium, followed by thorough mixing. The prepared drug-containing medium was then added to each well, resulting in a final volume of 4 mL per well for CCCP and oligomycin treatment, and 3ml for thapsigargin treatment. Final working concentrations were 10 µM CCCP, 1.25 ng/mL oligomycin, and 500 nM thapsigargin. Cells treated with CCCP or oligomycin were incubated for 16 h, whereas thapsigargin-treated cells were incubated for 6 h. Following treatment, cells were harvested for downstream analyses.

### Western blotting

HEK293T cells were lysed in 1× RIPA buffer prepared from 10× RIPA lysis buffer (MilliporeSigma, #20-188) supplemented with a protease inhibitor cocktail (Roche, #04693116001). Cell lysates were sonicated using a Bioruptor with cycles of 30 s on and 30 s off for 25 cycles, followed by centrifugation at 15,000 × g for 15 min at 4 °C. Total protein concentrations were determined using the Pierce 660 nm Protein Assay Reagent (Thermo Fisher Scientific, #22660). Protein samples were mixed with 4× Laemmli sample buffer and heated at 95 °C to denature proteins. Denatured samples were resolved by SDS–PAGE on homemade 12% Bis-Tris gels at 150 V for 1.5 h and transferred onto 0.45 µm nitrocellulose membranes (Cytiva, #10600002) at 200 mA for 2 h. Membranes were blocked with 5% non-fat dry milk (RPI, M17200-1000.0) at room temperature for 1 h. Primary antibodies were diluted in TBST buffer (25 mM Tris-HCl, pH 7.4, 150 mM NaCl, 1 percent BSA, and 0.1 percent Tween-20). The following primary antibodies were used: rabbit anti-ATF4 (Cell Signaling Technology, #11815, 1:1,000), mouse anti-β-actin (Proteintech, #66009-1-Ig, 1:5,000), rabbit anti-GFP (Cell Signaling Technology, #2956, 1:2,000), and rabbit anti-FLAG (Cell Signaling Technology, #14793, 1:2,000). Membranes were incubated with LI-COR infrared secondary antibodies, including goat anti-rabbit IgG IRDye 800CW (LI-COR: #926-32211, 1:20,000) and goat anti-mouse IgG IRDye 680RD (LI-COR #926-68070, 1:20,000). Blots were imaged using a ChemiDoc imaging system (Bio-Rad). Digital images were processed and quantified using ImageJ software.

### Co-immunoprecipitation

To analyze interactions between DELE1^CTD^ and designed binders, HEK293T cells were transiently co-transfected with 5 µg of FLAG-tagged DELE1^CTD^-GFP plasmid and 5 µg of GFP-tagged binder plasmids using Lipofectamine 3000 transfection reagent. A GFP empty vector was used as a negative control. Cells were harvested 48 h after transfection and lysed in lysis buffer containing 25 mM Tris-HCl (pH 7.5), 150 mM NaCl, 0.5 mM EDTA, 0.05 percent NP-40, 2.5% glycerol, and PMSF. A 50 µL aliquot of each lysate was mixed with 4× Laemmli sample buffer (Bio-Rad, #1610747) and heated at 95 °C for 5 min to serve as input controls. The remaining lysates were incubated with 20 µL of anti-FLAG M2 magnetic beads (Sigma-Aldrich, #M8823) for 3h at 4 °C with gentle rotation. Beads were washed three times with wash buffer containing 25 mM Tris-HCl, 150 mM NaCl, and 0.01% NP-40. Proteins bound to the magnetic beads were eluted by boiling in 2× Laemmli sample buffer (Bio-Rad, #1610737) at 95 °C for 5 min. Eluted samples and input controls were analyzed by SDS–PAGE and immunoblotting using antibodies described in the Western blotting section.

### Fluorescence microscopy

To examine how DELE1 binders influence mitochondrial elongation and network reformation following relief of mitochondrial stress, U2OS human osteosarcoma cells were used because of their relatively flat morphology, which facilitates high-quality live-cell imaging. Mitochondria were visualized by transient expression of the p00338 plasmid, which encodes pre-CoxIV fused to blue fluorescent protein. DELE1 binders were expressed using pXG289-Binder1-mClover or pXG289-Binder5-mClover plasmids, which encode Binder1 or Binder5 fused to mClover, enabling visualization and assessment of binder expression by green fluorescence.

For each independent experiment, 1 × 10^5^ passage 10 or passage 11 U2OS cells were seeded into 35 mm glass-bottom dishes in 3 mL of Dulbecco’s modified Eagle medium supplemented with 10% fetal bovine serum and 1% penicillin–streptomycin. Cells were allowed to adhere for 12 h prior to transfection. Transfections were performed using FuGENE 4K Transfection Reagent according to the manufacturer’s instructions with the following plasmid combinations: vehicle control, 0.25 µg p00338; Binder1 condition, 0.25 µg p00338 plus 0.50 µg pXG289-Binder1-mClover; and Binder5 condition, 0.25 µg p00338 plus 0.50 µg pXG289-Binder5-mClover. Three independent biological replicates were performed for each condition. After 40h of transient expression, cells were treated with 10 µM CCCP for 2h to induce mitochondrial stress. Following treatment, medium was aspirated, cells were washed once with 1 mL of Dulbecco’s phosphate-buffered saline, and 3 mL of pre-warmed fresh medium was added to initiate mitochondrial network recovery. Live-cell imaging was performed using an Axio Observer Z1 microscope equipped with a 61× oil-immersion objective. Images were acquired at baseline prior to CCCP treatment, after 2 h of treatment, and at 4 h, 8 h, and 24 h following medium replacement. Fluorescence excitation was provided by a PhotoFluor LM-75 light source, and images were captured using an AxioCam MRm camera with 358 nm and 488 nm excitation channels. Exposure times were 3000 ms for both channels. Image acquisition and export were performed using Zeiss Zen Pro software. For each independent experiment, 10 cells displaying clearly resolved mitochondrial morphology under blue fluorescent protein signal were randomly selected from distinct regions of the dish for analysis. Cells in the Binder1 and Binder5 conditions were required to be positive for both blue fluorescent protein and mClover fluorescence to ensure expression of both the mitochondrial marker and the corresponding DELE1 binder. The proportion of cells exhibiting fused mitochondrial networks was quantified for each condition and time point. Data analysis and plotting were performed using GraphPad Prism 10. Statistical significance was assessed using one-way ANOVA.

---

**Figure S1, related to Figure 1.**
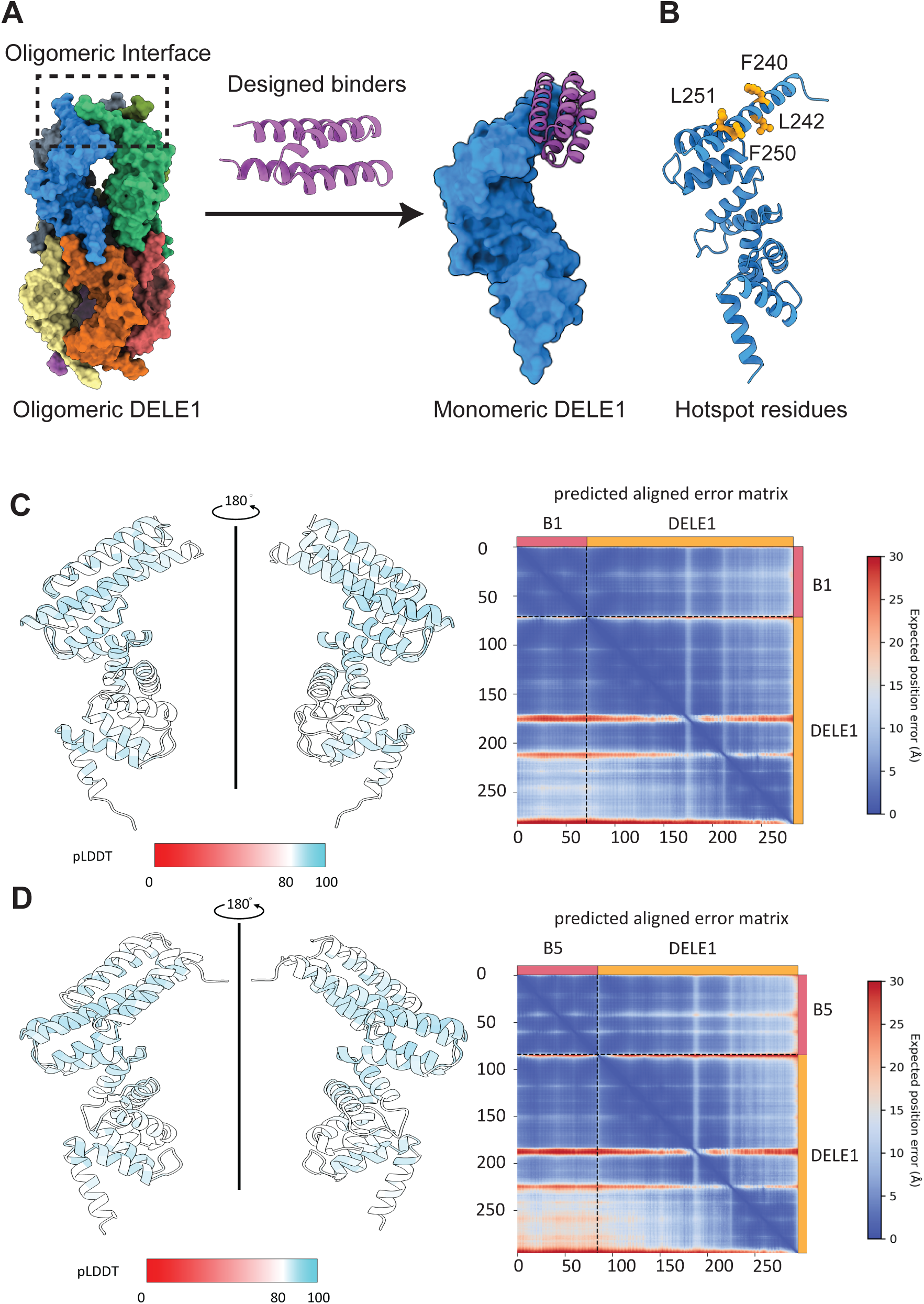
Structural rationale for targeting the DELE1 oligomerization interface using de novo protein design. (**A**) Surface representation of the oligomeric DELE1^CTD^ (PDB: 8D9X) assembly highlighting the oligomerization interface. Individual DELE1 subunits are colored distinctly to illustrate the multimeric architecture. Designed binders were engineered to engage the α1 helix at the oligomerization interface, thereby stabilizing a monomeric DELE1CTD species. (**B**) Ribbon representation of a monomeric DELE1^CTD^ subunit highlighting hydrophobic hotspot residues on the α1 helix (F240, L242, F250, and L251) selected for binder targeting. These residues form a contiguous interaction surface that is buried in the oligomeric assembly and were used to guide de novo binder design. (**C–D**) AlphaFold3-predicted structures of the DELE1–binder1 (C) and DELE1–binder5 (D) complexes. Left, structural models are shown in two orientations related by a 180° rotation. Structures are colored according to per-residue pLDDT confidence scores (scale shown below; red, low confidence; teal, high confidence). Right, PAE matrices for each complex. Low PAE values (blue) indicate high confidence in the relative positioning between residues, whereas higher PAE values (red) indicate increased uncertainty.

**Figure S2, related to Figure 2.**
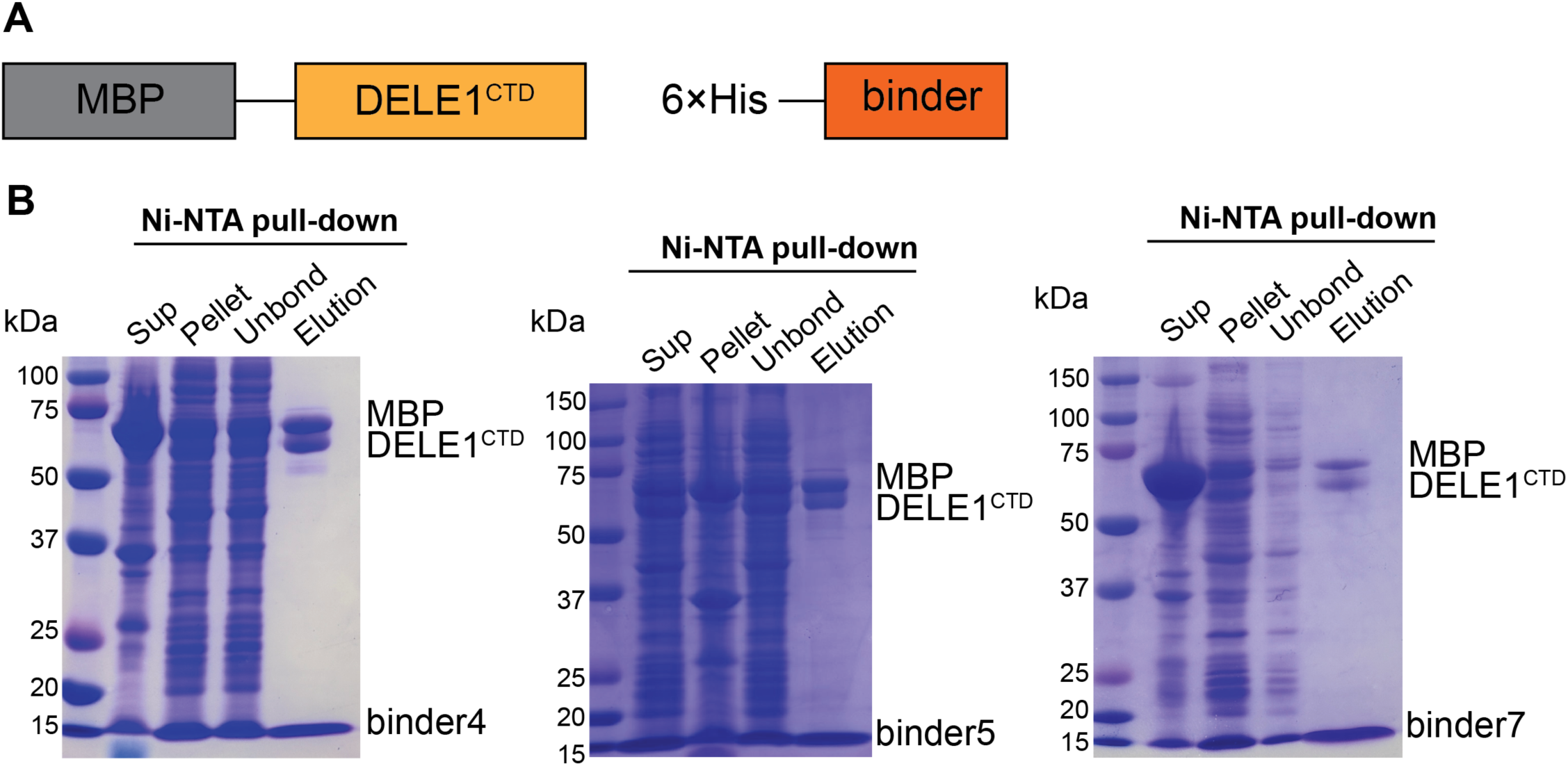
Co-expression and Ni–NTA pulldown of DELE1 with designed binders. (**A**) Schematic of the bacterial co-expression constructs. MBP-tagged DELE1^CTD^ was co-expressed with individual 6His-tagged binders, enabling affinity purification through the binder tag. (**B**) Coomassie-stained SDS–PAGE analysis of Ni–NTA pulldown experiments for representative binders (binder4, binder5, and binder7). Lanes show soluble lysate (Sup), insoluble fraction (Pellet), unbound material (Unbound), and eluted fractions (Elution). MBP-DELE1^CTD^ co-purifies with the corresponding 6His-tagged binders, indicating stable complex formation.

**Figure S3, related to Figure 2.**
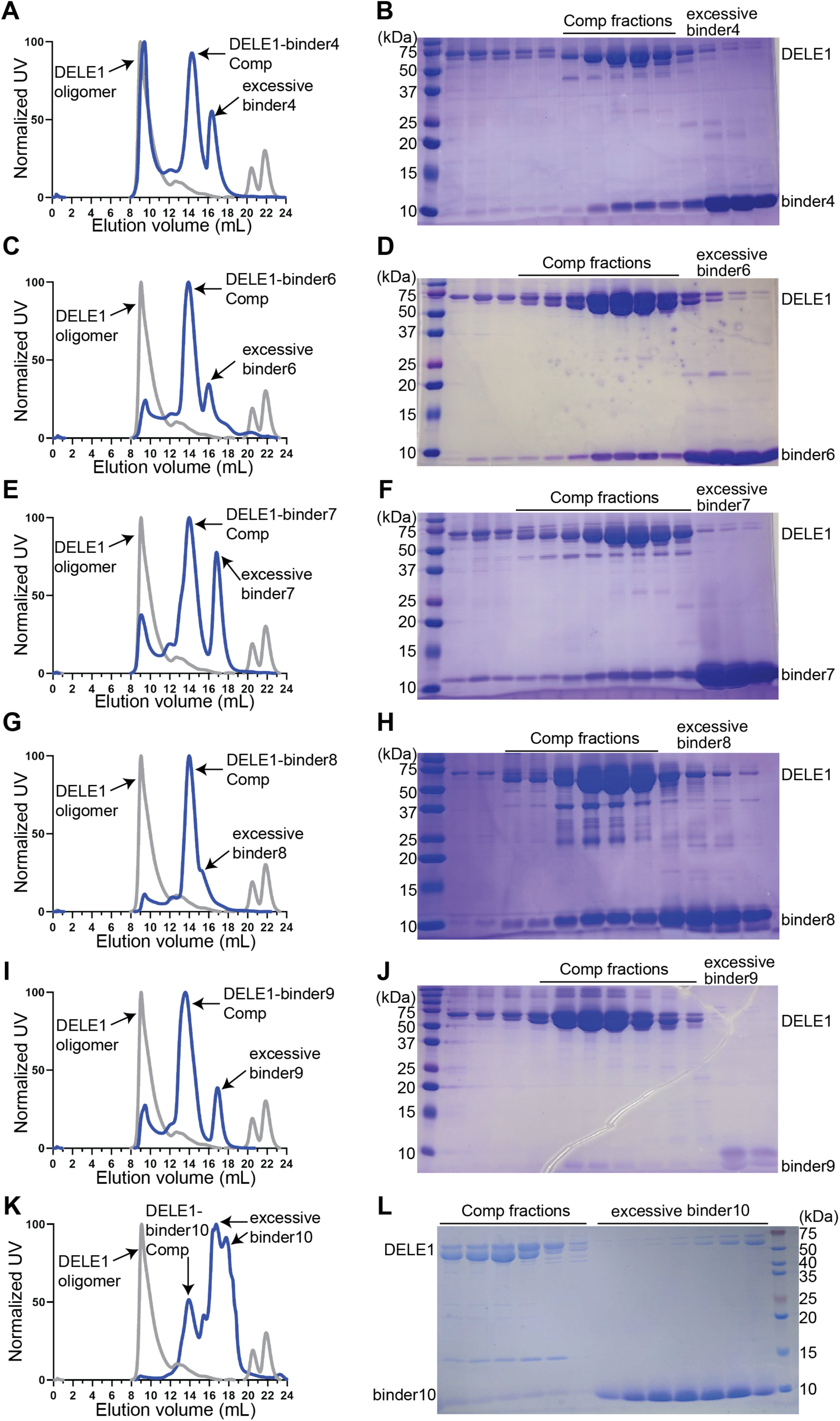
Additional designed binders form stable complexes with DELE1 in vitro. (**A, C, E, G, I, K**) Size-exclusion chromatography profiles of MBP-DELE1 co-expressed with additional 6His-tagged binders (binder4, binder6, binder7, binder8, binder9, and binder10). Elution profiles show the DELE1–binder complex (blue trace), oligomeric DELE1 species (gray trace), and excess unbound binder (black trace). Elution volumes are indicated on the x-axis. (**B, D, F, H, J, L**) Coomassie-stained SDS–PAGE analysis of SEC fractions corresponding to the major SEC peaks shown in panels A, C, E, G, I, and K. Fractions from the dominant peak contain both MBP-DELE1 and the corresponding binder, indicating complex formation. For binder10, co-elution with DELE1 was reduced relative to other binders, indicating a weaker complex formation. Lanes labeled “excess binder” contain unbound binder carried through affinity purification. These data extend the results shown in Figure 2 and demonstrate that multiple independently designed binders associate with DELE1 and form homogeneous complexes in vitro.

**Figure S4, related to Figure 3.**
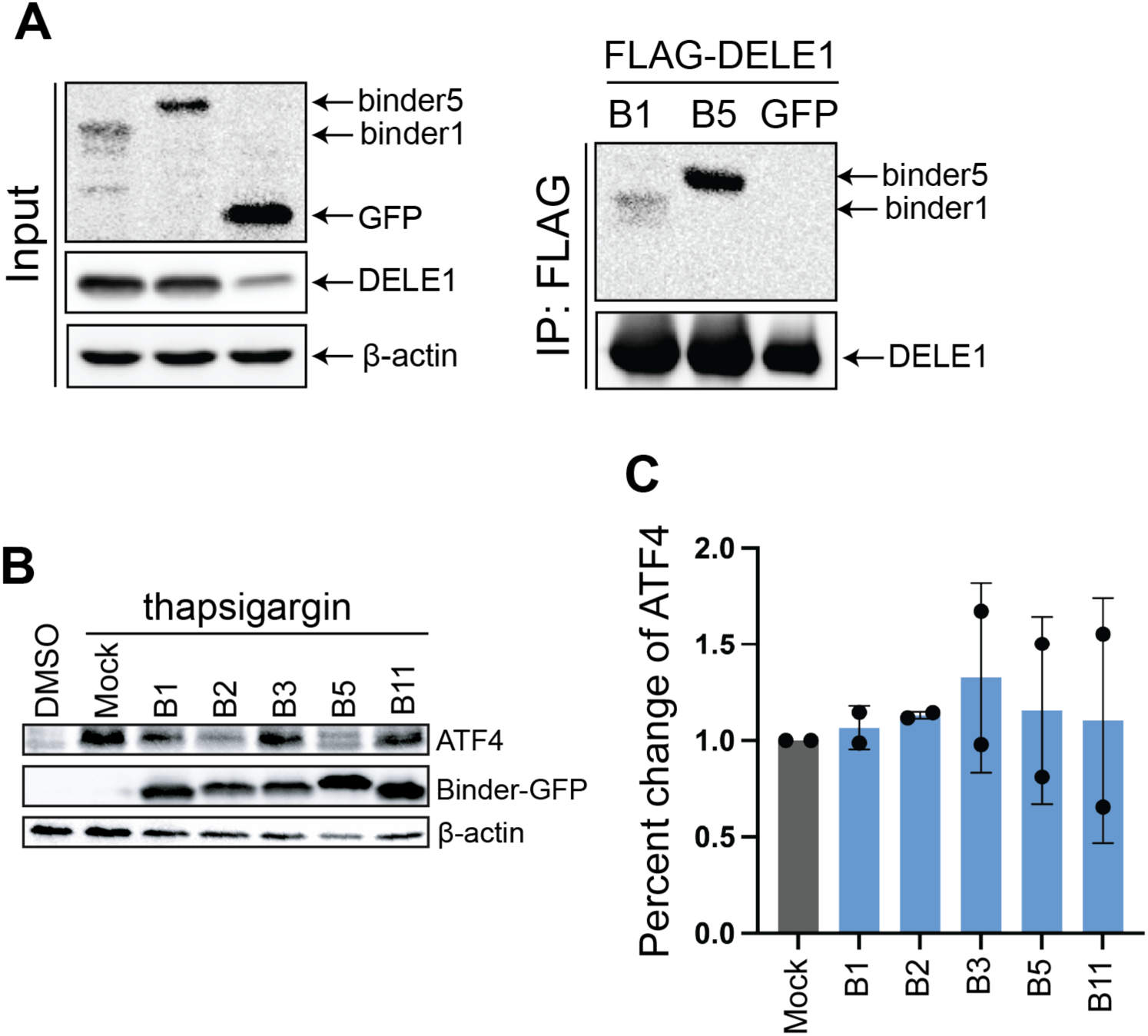
Designed binders directly interact with DELE1 in cells and do not suppress endoplasmic reticulum stress induced ATF4. (**A**) Co-immunoprecipitation analysis of binder association with DELE1 in HEK293T cells. FLAG-tagged DELE1-GFP was co-expressed with GFP, binder1-GFP, or binder5-GFP. Whole-cell lysates (Input) and anti-FLAG immunoprecipitants (IP: FLAG) were analyzed by immunoblotting for DELE1 and GFP-tagged binders. (**B**) Immunoblot analysis of ATF4 levels in HEK293T cells expressing GFP-tagged binders and treated with thapsigargin for 6 hrs. Binder expression was monitored by GFP immunoblotting, and β-actin served as a loading control. (C) Quantification of ATF4 levels from CCCP-treated samples shown in panel **B**, normalized to β-actin and expressed relative to mock-transfected cells. Data represent mean ± s.d. from 2 independent experiments. DMSO represents cells treated with DMSO and serves as the blank control, rather than treatment with CCCP or oligomycin. Mock represents cells treated with CCCP or oligomycin as a positive control; however, no binders were transfected or overexpressed in these cells.

**Figure S5, related to Figure 4.**
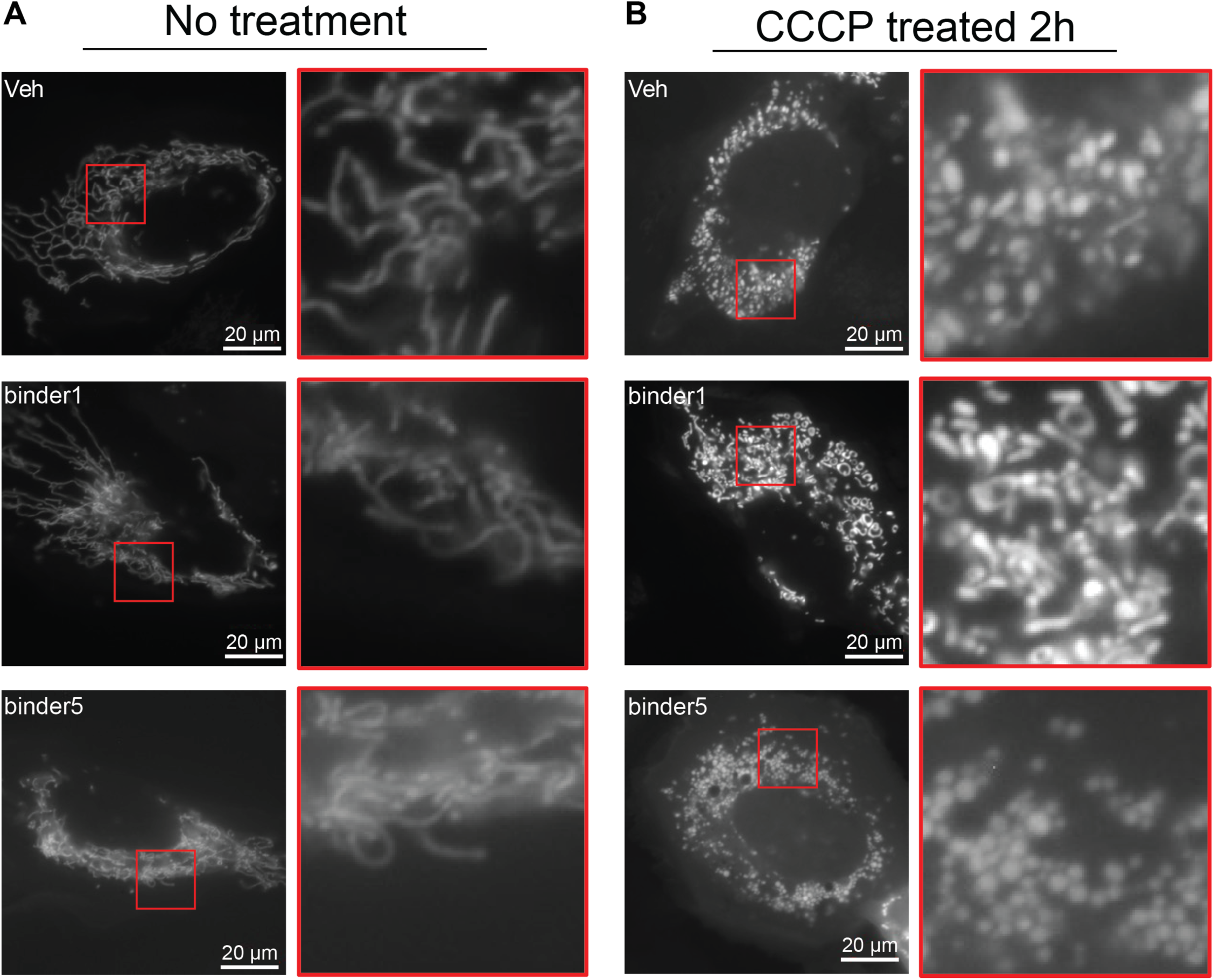
Mitochondrial stress triggers fragmented network morphology. Representative fluorescence microscopy images of mitochondrial morphology in U2OS cells before and after CCCP treatment. Cells were transiently transfected with BFP-tagged pre-CoxIV to label mitochondria and, where indicated, co-expressed with binder1–GFP or binder5–GFP. (**A**) Under untreated conditions, control cells and cells expressing binder1 or binder5 display an interconnected, elongated mitochondrial network. (**B**) Following 2 hours of CCCP treatment, mitochondria undergo robust fragmentation in all conditions, regardless of binder expression. For each condition, a higher-magnification view of the boxed region (red) is shown to the right. Scale bars, 20 µm. Veh represents cells treated with as a positive control; however, no binders were transfected or overexpressed in these cells. **: <0.01, *:<0.05, ns: not significant.

**Figure S6, related to Figure 5.**
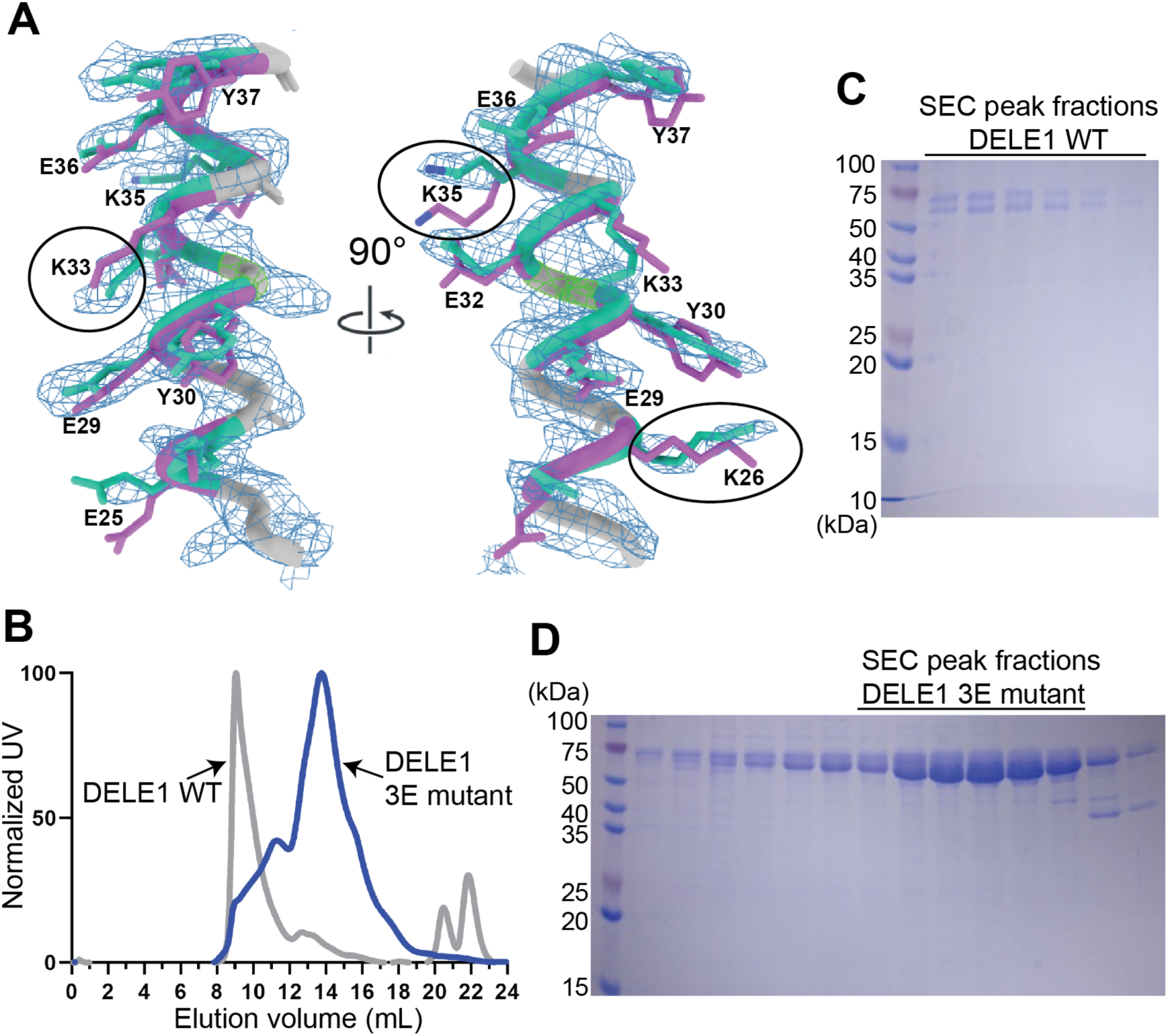
Structural validation of binder5 and biochemical characterization of the DELE1 3E mutant. (A) Representative electron density map (2Fo–Fc) for binder5 contoured around the refined atomic model, shown in two orientations related by a 90° rotation. Side chains of selected residues are labeled to illustrate the quality of density. Side-chain conformations from the crystal structure (teal) are overlaid with those from the design model (purple) to highlight local differences between the experimental structure and the predicted model. (B) SEC profiles of MBP-DELE1 wild-type (gray trace) and the L229E/F240E/F250E triple mutant (“3E”, blue trace). The 3E mutant elutes at a later volume relative to wild-type DELE1, indicating a shift toward a smaller apparent molecular weight species. (C) Coomassie-stained SDS–PAGE analysis of SEC peak fractions for MBP-DELE1 wild-type. (D) Coomassie-stained SDS–PAGE analysis of SEC peak fractions for the MBP-DELE1 3E mutant.

**Figure S7, related toc Figure 6.**
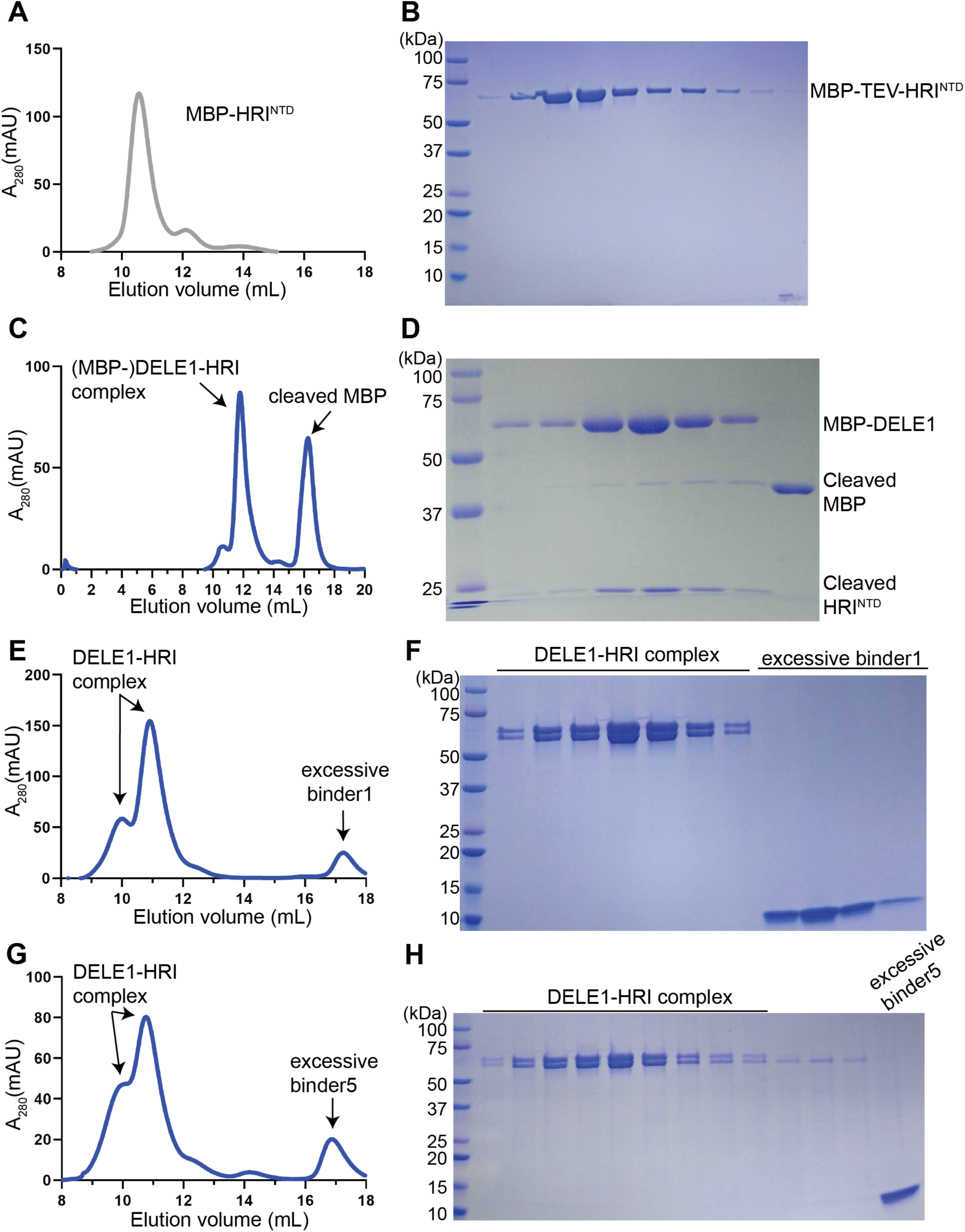
Direct interaction between DELE1 and HRI-NTD is preserved in the presence of designed binders. (**A**) SEC profile of purified MBP–TEV–HRI^NTD^, showing a homogeneous, early-eluting peak. (**B**) Coomassie-stained SDS–PAGE analysis of SEC fractions from panel A, confirming purity of MBP–TEV-HRI^NTD^. (**C**) SEC profile of the DELE1–HRI complex following TEV protease treatment. Two major high–molecular weight species are observed, corresponding to complexes containing MBP–DELE1 and cleaved HRI^NTD^. (**D**) SDS–PAGE analysis of SEC fractions from panel C, showing co-elution of MBP–DELE1 and cleaved HRI^NTD^. The cleaved MBP was eluted in later fractions. (**E**) SEC analysis of the pre-formed DELE1–HRI complex incubated with excess binder1. The DELE1–HRI complex remains intact, while excess binder1 elutes as a separate low–molecular weight peak. (**F**) SDS–PAGE analysis of SEC fractions from panel E, confirming retention of the DELE1–HRI complex and absence of binder1 in the complex fractions. (**G**) SEC analysis of the pre-formed DELE1–HRI complex incubated with excess binder5, showing a stable DELE1–HRI complex and a separate low–molecular weight peak corresponding to excess binder5. (H) SDS–PAGE analysis of SEC fractions from panel G, confirming that binder5 does not disrupt the DELE1–HRI complex.

